# Differentiation of ciliated human midbrain-derived LUHMES neurons

**DOI:** 10.1101/2020.06.04.134478

**Authors:** Gilbert Lauter, Andrea Coschiera, Masahito Yoshihara, Debora Sugiaman-Trapman, Sini Ezer, Shalini Sethurathinam, Shintaro Katayama, Juha Kere, Peter Swoboda

**Author notes:** Correspondence: Peter Swoboda, Karolinska Institute, Department of Biosciences and Nutrition, Campus Flemingsberg - NEO Building, Hälsovägen 9, SE-141 83 Huddinge, Sweden. equal contribution – joint first authorship (Gilbert Lauter, Andrea Coschiera, Masahito Yoshihara). equal contribution – joint last authorship: JK, PS.

## Abstract

Many human cell types are ciliated, including neural progenitors and differentiated neurons. Ciliopathies are characterized by defective cilia and comprise various disease states, including brain phenotypes, where the underlying biological pathways are largely unknown. Our understanding of neuronal cilia is rudimentary and an easy-to-maintain, ciliated human neuronal cell model is missing.

LUHMES is a ciliated neuronal cell line derived from human fetal mesencephalon. LUHMES cells can easily be maintained and differentiated into mature, functional neurons within one week. They have a single primary cilium as proliferating progenitor cells and as post-mitotic, differentiating neurons. These developmental stages are completely separable within one day of culture condition change. The Sonic Hedgehog (SHH) signaling pathway is active in differentiating LUHMES neurons. RNA-seq time course analyses reveal molecular pathways and gene-regulatory networks critical for ciliogenesis and axon outgrowth at the interface between progenitor cell proliferation, polarization and neuronal differentiation. Gene expression dynamics of cultured LUHMES neurons faithfully mimic the corresponding in vivo dynamics of human fetal midbrain.

In LUHMES, neuronal cilia biology can be investigated along a complete timeline: from proliferation through differentiation to mature neurons.

**Summary Statement:** With LUHMES, a ciliated human neuronal cell model, the underlying “neurobiology” of cilia and ciliopathies can be investigated along a complete time line: from proliferation through differentiation to mature neurons.

## Introduction

Primary cilia are non-motile, hair-like cell surface protrusions present in single copy on polarized cell types. The cilium consists of a microtubular core structure, the axoneme, surrounded by a specialized membrane. The axoneme is a continuous outgrowth from the centriole-derived basal body, which anchors the ciliary shaft within the cytoplasm of the cell. A transition zone at the base of the axoneme forms a ciliary gate where cargo is sorted for import and export to and from the ciliary compartment. The maintenance of cilia depends on selective transport of protein components mediated by the intraflagellar transport (IFT) machinery (Ishikawa and Wallace, 2011).

Primary cilia are not restricted to terminally differentiated cell types, but can also be found on proliferating cells, like neural progenitors. The centriole serves in two mutually exclusive functions: either as basal body and axonemal template in ciliogenesis or as microtubule-organizing center (MTOC) of the centrosome. Cilia are thus tightly associated with the dynamics of the cell cycle (Malicki and Johnson, 2017). Ciliogenesis is linked to the G0/G1 phase of the cell cycle. Prior to cell division the cilium is resorbed to free a pair of centrioles from the ciliary base. During S and G2 phases centrioles are duplicated and mature into functional centrosomes, which organize the formation of the bipolar spindle for sister chromatid separation in M phase (Pletz et al., 2013; Ford et al., 2018; Joukov and De Nicolo, 2019).

Primary cilia are antennae that receive and transduce signals from the immediate environment of a cell. The primary cilium as a signaling hub integrates several aspects of the SHH pathway, but is also involved in other signal transduction pathways, such as WNT, Notch, platelet-derived growth factor (PDGF) or multiple G-protein coupled receptors (Goetz and Anderson, 2010; Lauter et al., 2018; Liu et al., 2018; Wheway et al., 2018). In addition to being a receiver, the primary cilium has also gained attention as a source of signals to its cellular environment (Garcia et al., 2018; Nachury and Mick, 2019).

Cilia are highly conserved within the eukaryotic kingdom (Piasecki et al., 2010) and in humans they can be found on many different cell types (Reiter and Leroux, 2017). Primary cilia are a common feature of polarized cells, and they are abundant in the brain, both during development and at mature stages, extending from progenitor cells as well as from terminally differentiated neurons and astrocytes (Bishop et al., 2007; Arellano et al., 2012; Sarkisian and Guadiana, 2015). Disruption in the structure, maintenance or function of primary cilia gives rise to a group of highly divergent disorders collectively termed ciliopathies.

Due to the widespread distribution of cilia, ciliopathies display pleiotropic features, including brain phenotypes (Mitchison and Valente, 2017; Reiter and Leroux, 2017). A number of neurodevelopmental disruptions that result from SHH-dependent patterning defects and certain aspects of neuropsychiatric dysfunctions, as observed in Joubert Syndrome and autism-spectrum disorders, have been connected to ciliary aberrations (Massinen et al., 2011; Lauter et al., 2018; Youn and Han, 2018; Thomas et al., 2019). The associations between changes in ciliary structure and function and the manifestation of a particular (brain) disorder vary in solidity and strength. The causality of the link is also a matter of debate. Thus, investigations of the underlying molecular and cellular mechanisms, possibly cilia-associated, are urgently needed.

Currently, multiple mammalian cell culture models exist for the investigation of cilia biology, including ciliary aspects of sensory perception and signal transduction. Despite the undisputed value of existing models, no easily amenable human neuronal cell model exists, which would allow studying ciliogenesis, cilia structure and function in (neural) progenitor cells during all stages of the cell cycle, as well as during development and neuronal differentiation. To fill this gap, we set up the **Lu**nd **h**uman **mes**encephalic (LUHMES) cell line as a model for studying primary cilia in human neurons. Ciliated LUHMES cells are easy to maintain and can be differentiated into fully functional neurons within about one week.

## Results

### The human midbrain-derived LUHMES cell model is ciliated

The LUHMES cell line is an easy-to-maintain human neuronal cell model and candidate for studying cilia biology (Scholz et al., 2011). We used immunocytochemistry to identify neuronal cilia during both cell proliferation and after various days past the induction of differentiation into functional neurons. ADP Ribosylation Factor Like GTPase 13B (ARL13B) is a widely used marker for cilia that is specifically and strongly enriched in the cilium (Caspary et al., 2007, 2016; Ferent et al., 2019). Immunofluorescence stainings revealed that ARL13B is expressed and specifically localized in LUHMES cells at the stage of cell proliferation (d0) as well as during differentiation into neurons (d1 to d6) (**Figure 1A**), suggesting that LUHMES cells and neurons are ciliated. To confirm ciliary localization we looked at double stainings of ARL13B together with other known ciliary markers. Pericentrin (PCNT) localizes to centrioles and to centriole-derived ciliary basal bodies (Pitaval et al., 2010; Ishikawa and Marshall, 2011; Pletz et al., 2013). ARL13B and PCNT co-localized to the ciliary shaft and basal body, respectively (**Figure 1B**), corroborating the presence of cilia. Proper structure and function of cilia depend on intraflagellar transport (IFT) (Ishikawa and Marshall, 2011). As expected, IFT88 protein co-localized together with ARL13B along the ciliary shaft (**Figure 1C**), further supporting the presence of neuronal cilia. Other ciliary markers, such as detyrosinated tubulin and RAB8, also showed expression and specific localization to cilia (data not shown). Using immunofluorescence stainings we could thus demonstrate the presence of cilia on LUHMES cells and neurons.

**Figure 1:**
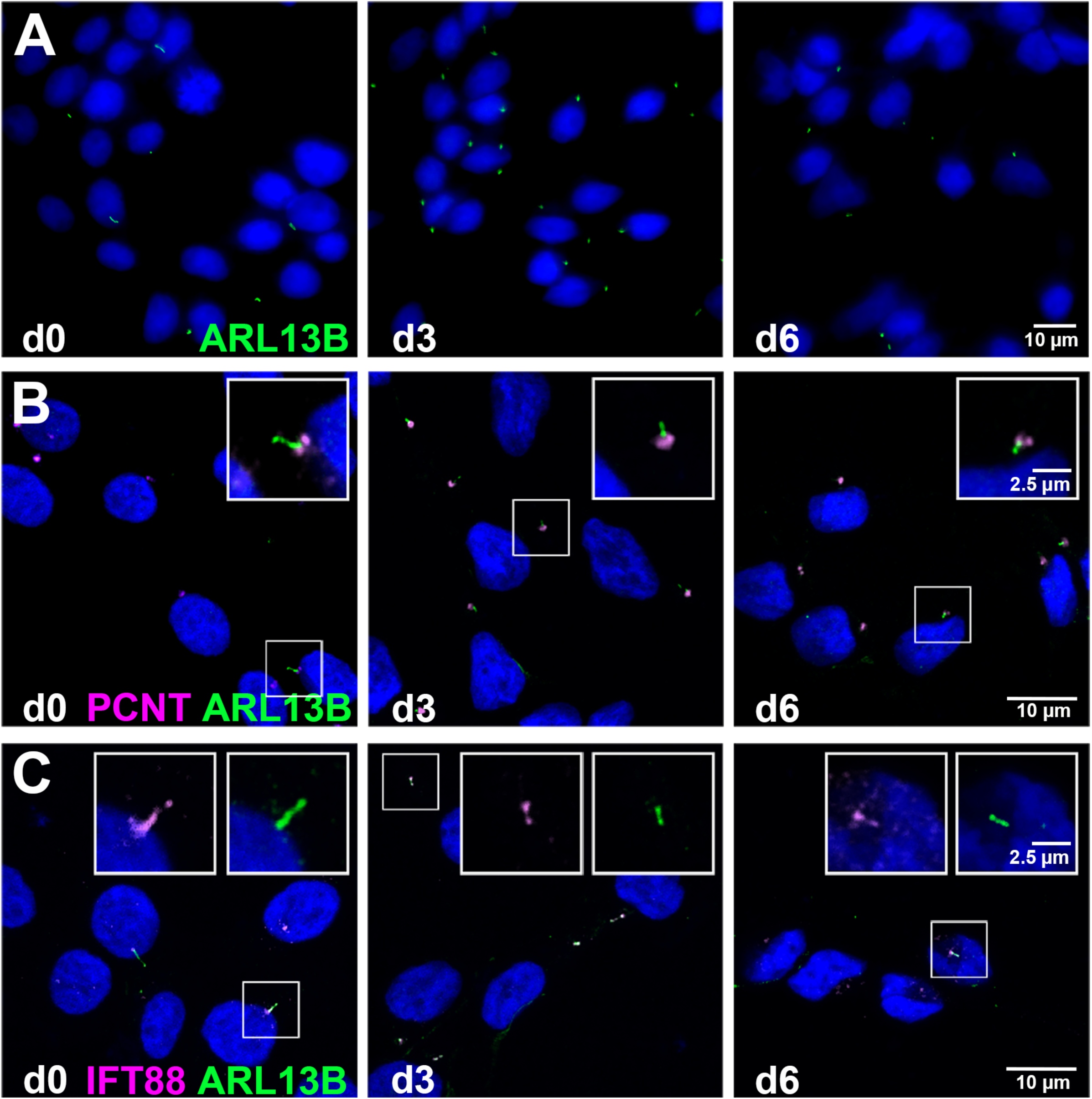
Human midbrain-derived LUHMES neurons are ciliated. Ciliary structures are detectable on LUHMES cells during proliferation (d0) and at various time points after induction of neuronal differentiation (d3 and d6). Immunofluorescence stainings of known markers for cilia: **(A)** ARL13B (green) marks the ciliary shaft and is detectable at all stages analyzed. **(B)** ARL13B (green) consistently co-localizes with the basal body marker pericentrin (PCNT) (magenta). **(C)** IFT88 protein (magenta), important for intraflagellar transport (IFT) along the ciliary axoneme, is co-expressed with ARL13B (green) at all stages analyzed. The boxed areas indicate magnified regions. In all panels the scale bar represents 10 µm, in the boxed areas 2.5 µm.

To unambiguously confirm the presence and the nature of cilia on LUHMES cells and neurons, we performed transmission electron microscopy (TEM). Using TEM we could clearly verify the presence of extended cilia (**Figure 2**). We detected all the defining parts of cilia in cross-as well as longitudinal sections through the structure. Basal bodies displayed the characteristic configuration of nine microtubule triplets (**Figure 2A-B**), whereas more distal parts, including the ciliary axoneme, consisted of the typical microtubule doublet formation (**Figure 2C**) (Pletz et al., 2013). Basal body appendages and transition zone structures (**Figure 2B-E**) as well as a pronounced ciliary pocket were clearly distinguishable (**Figure 2C-G**). Of note, the apparent lack of dynein arms between the microtubule doublets (**Figure 2C**) and the arrangement of these microtubule doublets in a 9+0 configuration confirmed that LUHMES cilia are non-motile, primary cilia.

**Figure 2:**
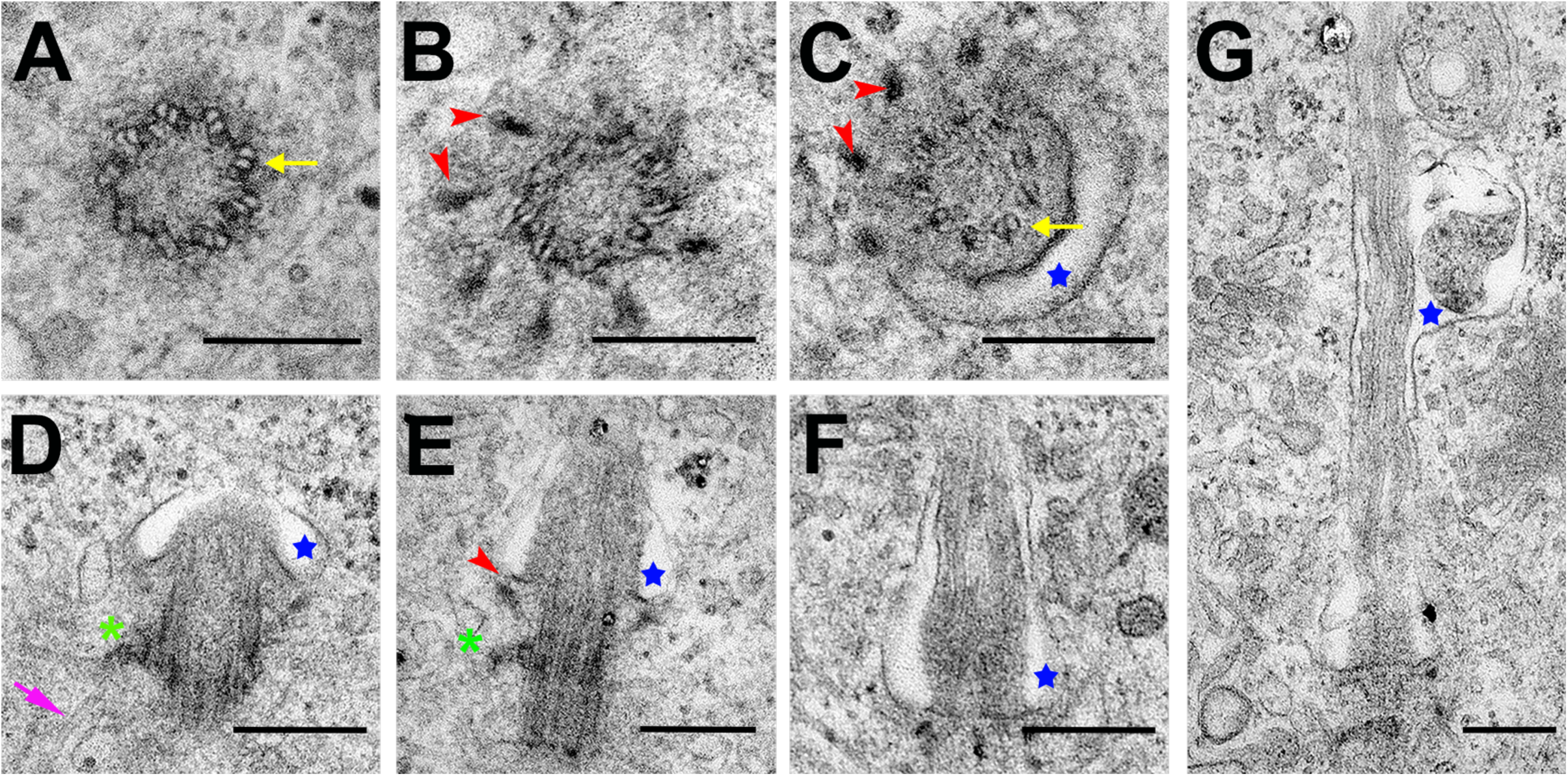
Cilia of differentiating LUHMES neurons (d3) in transmission electron microscopy. **(A)** A cross-section through the basal body reveals the nine-fold configuration of microtubule triplets (arrow). **(B)** A cross-section through the distal part of the basal body shows apparent transition fibers extending outward (arrowhead). **(C)** A slightly tilted cross-section reveals transition fibers (arrowheads – upper left) with the ciliary axoneme in the 9+0 configuration typical for primary cilia (microtubule doublets; arrow – lower right) and an adjacent ciliary pocket (blue star). **(D-G)** Longitudinal sections show cilia at different stages and angles (blue stars mark the ciliary pocket; arrowheads and green asterisks mark distal and subdistal appendages, respectively; the arrow in magenta indicates microtubules converging on subdistal appendages). In all panels the scale bar represents 250 nm.

Primary cilia are involved in the transduction of a variety of signaling pathways, including in vertebrates the coordination of the canonical Sonic Hedgehog (SHH) signaling pathway (Anvarian et al., 2019). To determine whether the primary cilia in human LUHMES neurons are functional signaling units, we examined the output of SHH signaling. We treated differentiating (d2) LUHMES neurons for 24 h with a SHH pathway activator, Smoothened agonist (SAG), and an antagonist, Cyclopamine, that modulate the canonical SHH pathway through the GLI transcription factors (Stanton et al., 2010). After treatment (on d3), we evaluated by qRT-PCR the gene expression of three downstream SHH pathway target genes: GLI1, HHIP and PTCH1. Samples treated with the SHH pathway activator SAG (500 nM) showed a significant upregulation of target gene expression, as compared to the samples treated with the antagonist Cyclopamine (10 µM) or the control samples (0.25% DMSO) (**Figure 3**). These observations confirm that the primary cilia in LUHMES transmit molecular signals, exemplified by the canonical SHH signal transduction pathway.

**Figure 3:**
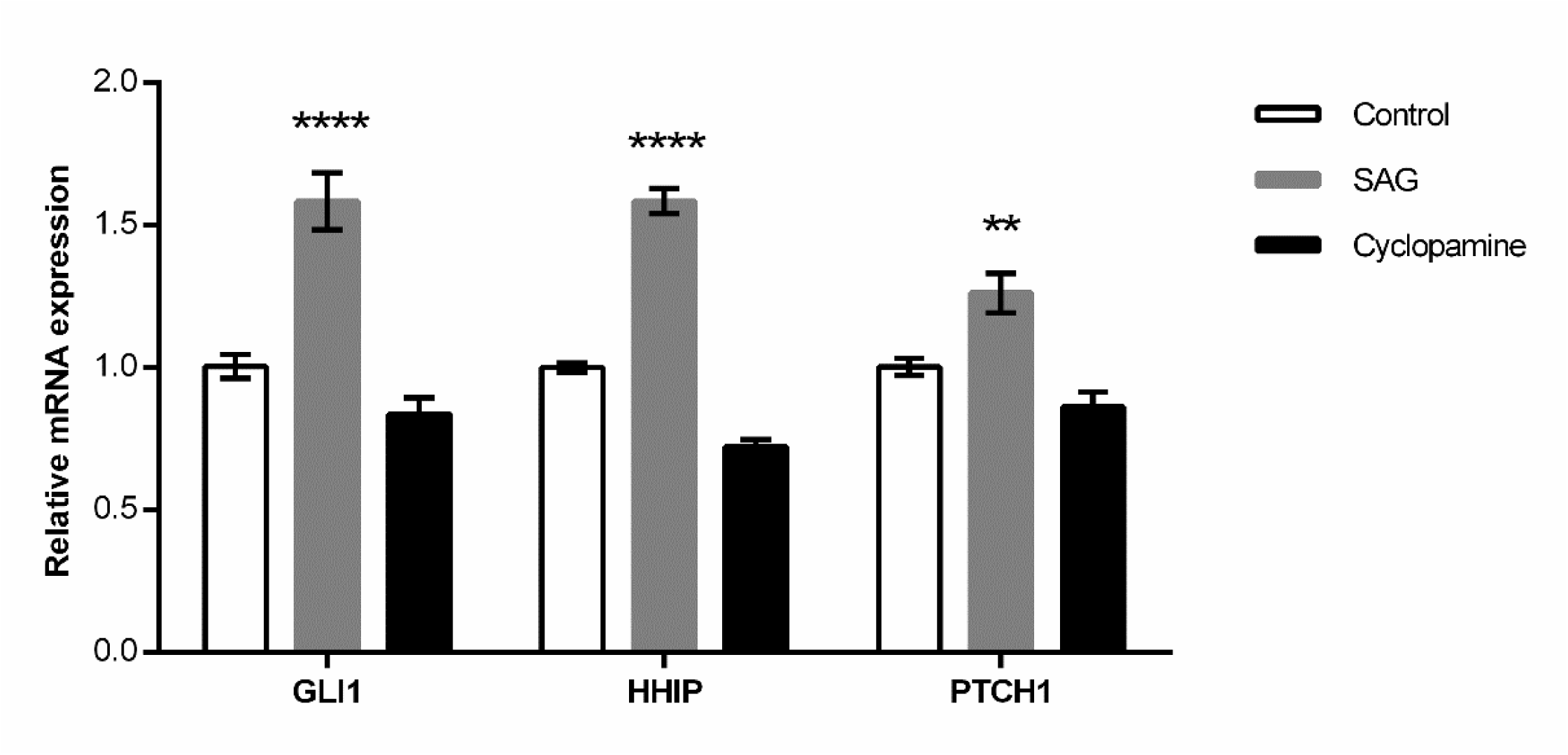
The ciliary Sonic Hedgehog (SHH) signaling pathway is active in differentiating, ciliated LUHMES neurons. When treating LUHMES neurons with a SHH pathway activator, Smoothened Agonist (SAG, 500 nM, 24h), the relative expression of SHH pathway target genes (GLI1, HHIP, PTCH1) is significantly upregulated as compared to the control (DMSO, 0.25%, 24h). Treatment with Cyclopamine (10 μM, 24h), a known SHH pathway antagonist, completely reverts this up-regulation. Statistical analysis: We conducted a regular two-way ANOVA (not repeated measures) with multiple comparisons between conditions. Mean values +/- SEM were normalized to GAPDH. The results are from two independent experiments with a total of four technical replicates. **: p < 0.005; ****: p < 0.0001.

### LUHMES cells and neurons are ciliated during both proliferation and differentiation

To establish the LUHMES cell model as a useful tool for studying human, neuronal cilia we adopted a simplified protocol for differentiating LUHMES cells into functional neurons: within about one week (d0 – d6) proliferating LUHMES cells can be differentiated into a state at which different neuronal differentiation and maturation markers are readily detectable and even solid electrophysiological assays are possible (Scholz et al., 2011). To better mimic in situ neuronal development, we used unsynchronized culture conditions for neuronal differentiation.

By following representative groups of several hundreds of cells through image capture and by manual counting, we determined that upon induction of neuronal differentiation at d0, LUHMES cells divide only once from d0 to d1 and then become post-mitotic (**Figure 4A-D**). The complete arrest of all cell proliferation after about one day post-induction of neuronal differentiation confirmed that the applied culturing protocol pushed all the cells into full differentiation mode. We did not observe any significant cell death (tested up to d6), as cell viability was consistently > 95% (**Figure 4E**). These features suggest that the adopted culture conditions (Scholz et al., 2011) were in the optimal range.

**Figure 4:**
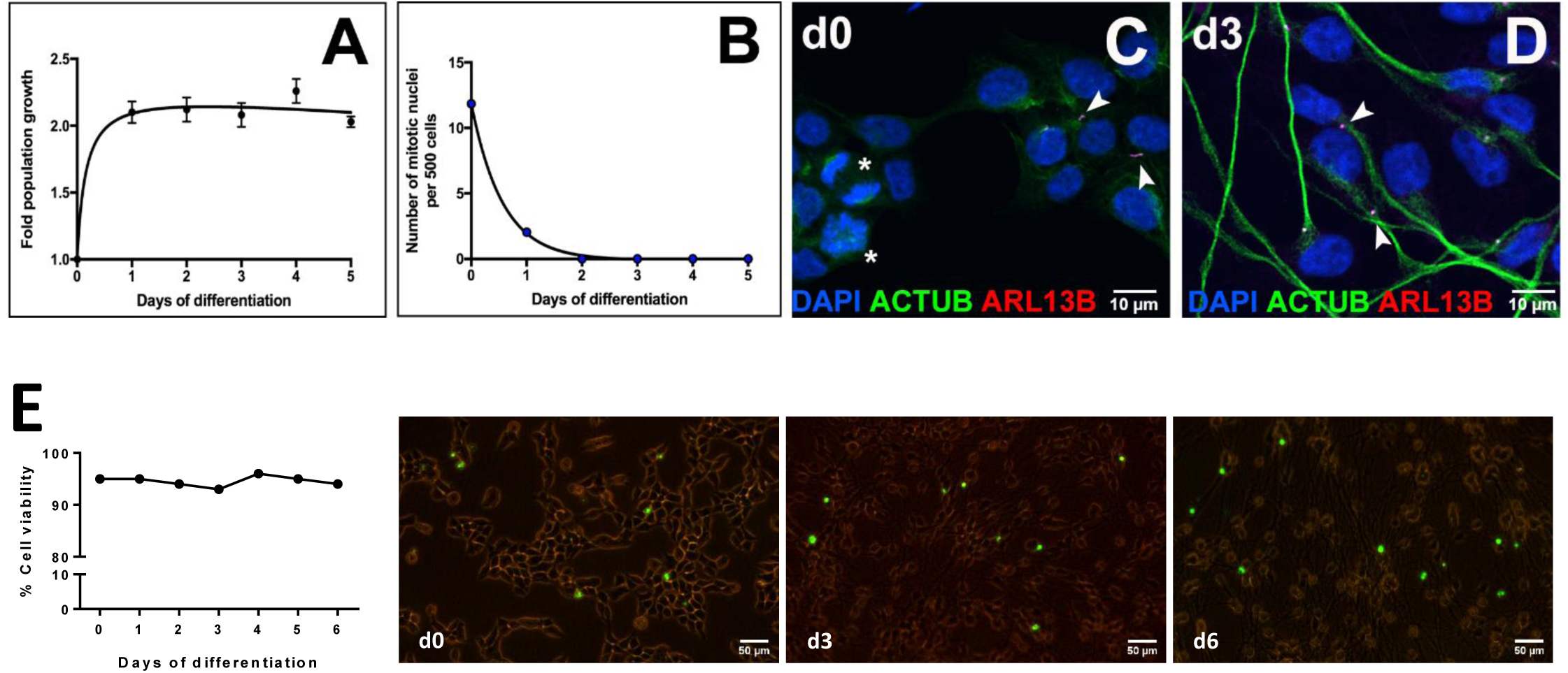
The differentiation of LUHMES cells into neurons is preceded by a complete, population-wide cell cycle arrest. A differentiating cell population culture was followed under the microscope from d0 to d6. **(A)** The cell population doubles between d0 and d1 and then remains constant, suggesting that only one cell division occurs during the first day after the induction of differentiation. **(B)** The number of mitotic cell nuclei was counted in a population of >500 cells. The presence of mitotic cell nuclei decreases rapidly and is undetectable at d2 after the induction of differentiation. **(C-D)** Immunofluorescence staining of nuclei (DAPI, blue), cilia (ARL13B, red) and neurites (ACTUB, green) at d0 and d3 of differentiation. Asterisks indicate mitotic nuclei and arrowheads indicate primary cilia. In both panels the scale bar represents 10 µm. **(E)** Cell and neuron viability in culture remains constant > 95%: representative fluorescence images from LUHMES cells and neurons at d0, d3 and d6 of differentiation indicate rare dead cells (marked with NucGreen Dead 488 Ready Probe). In all fluorescence microscopy panels the scale bar represents 50 µm.

To confirm that LUHMES cells develop cilia as they become post-mitotic and differentiate, we followed the expression of both ciliary (ARL13B) and neuronal differentiation markers. From loosely aggregated colonies of proliferating LUHMES cells at d0 (**Figure 5A**), differentiating neurons dispersed and progressively grew neurite extensions to eventually form a well-developed network of neurites, easily discernable on d5 or d6 (**Figure 5**). At any given time point during this developmental time course LUHMES cells and neurons display primary cilia (d0 to d6) (**Figures 1, 4, 5**). The percentage of ciliated cells and neurons was not uniform between stages. Proliferating LUHMES cells progressing through the cell cycle in our unsynchronized culture conditions displayed a much lower percentage of ciliation as compared to after the induction of (post-mitotic) neuronal differentiation. Ciliation of these neuronal populations quickly rose to well above 50%, reaching a plateau between d3 to d5 of differentiation (data not shown). This plateau of ciliation coincided well with the differentiating neuron population attaining a complex network of neurites during the same time period (d3 to d5).

**Figure 5:**
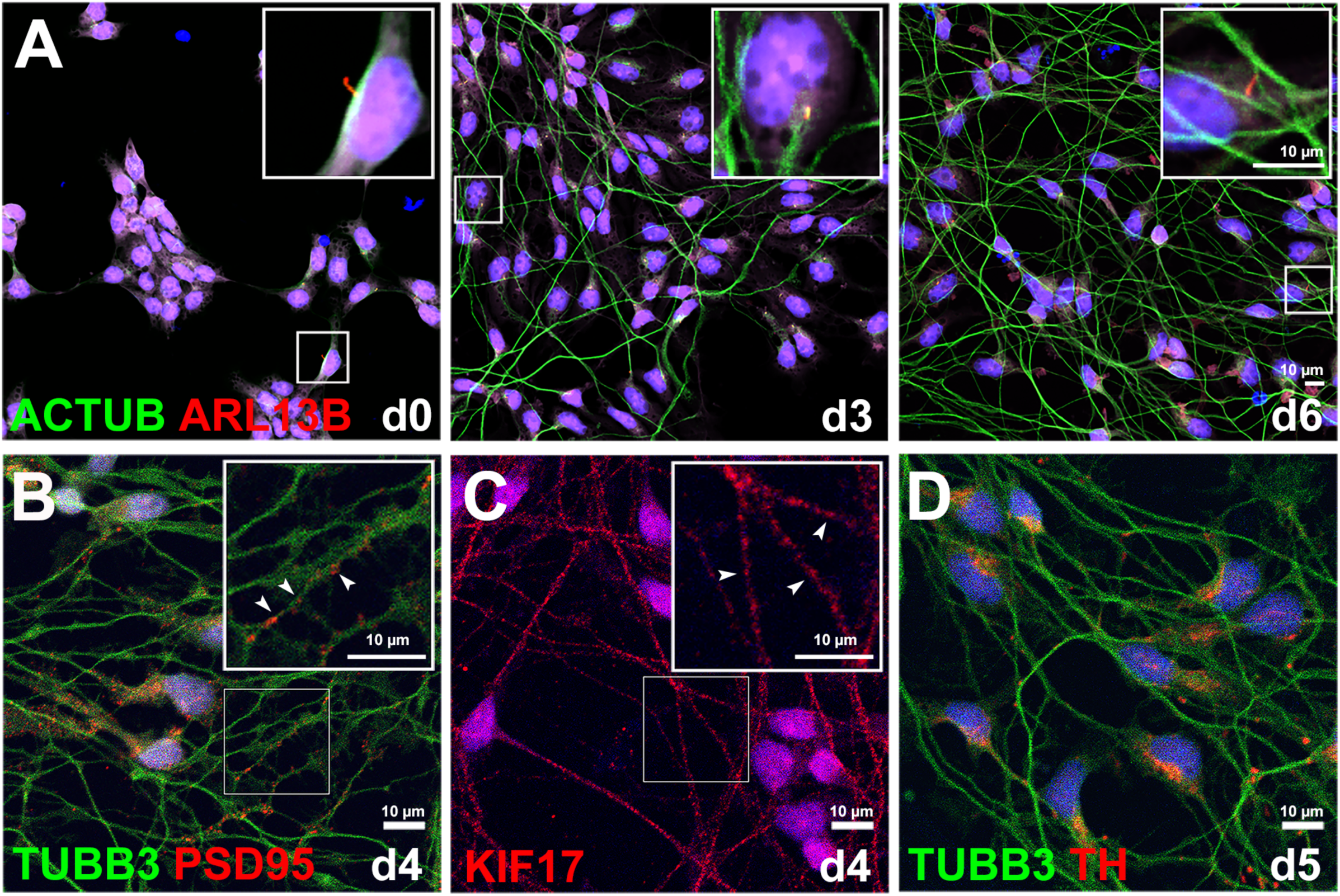
Cilia are present throughout the differentiation of LUHMES cells into neurons. Cell differentiation was followed using known markers for neuronal differentiation and cilia. Nuclei were stained with DAPI (blue). **(A)** Immunofluorescence staining of cilia (ARL13B, red) and developing neurites (ACTUB, green) at d0, d3 and d6 of differentiation. At d0 cells frequently cluster together and no neurite network is present. At d3 of differentiation neurons have started forming a neurite network expressing ACTUB. At d6 of differentiation neurons are highly interconnected and integrated into a neurite network. At all stages, primary cilia (inset) are detectable by ARL13B (red). **(B)** Along neurites (marked by TUBB3) clusters of PSD95, a post-synaptic marker, become visible at around d4 of differentiation. **(C)** The motor protein KIF17, a neurite transport marker, shows a characteristic punctate pattern at around d4 of differentiation. **(D)** Expression of the dopaminergic marker TH is detectable in the cell body at around d5 of differentiation. In panels with boxed areas these indicate the magnified regions shown as an inset in the upper right corner. In all panels the scale bar represents 10 µm.

LUHMES differentiation into functional neurons has been described elsewhere and our work faithfully replicated the findings, including the overall time frame (Scholz et al., 2011; Shah et al., 2016; Harischandra et al., 2020). Accordingly, we observed that, in addition to the ciliary marker ARL13B (**Figure 5A**), differentiating LUHMES neurons expressed the general neurite marker tubulin beta 3 (TUBB3) and the synaptic marker post-synaptic density protein 95 (PSD95) (**Figure 5B**), the neuronal transport marker kinesin 17 (KIF17) (**Figure 5C**) as well as tyrosine hydroxylase (TH), the terminal differentiation marker for dopaminergic neurons (**Figure 5D**) (Wulle and Schnitzler, 1989; Lee et al., 1990; Hunt et al., 1996; Franker et al., 2016). The expression of various markers for neuronal differentiation, maturation and function, suggested that our culture conditions successfully induced neuronal differentiation, in accordance with previous findings.

Proliferating, progenitor-type LUHMES cells (d0) differ strongly from differentiating LUHMES neurons (d1 to d6) (Scholz et al., 2011) with regard to anatomy and morphology, physiology and function, and a number of molecular aspects. One main difference is the absence of developing or differentiated neurites (dendrites and axons) in proliferating LUHMES cells. Despite these differences the position and localization of the primary cilium remained constant during both proliferation and differentiation (**Supplementary Figure S1**). Using regular monolayer-type culture conditions and immunofluorescence stainings (ARL13B) we found by visual inspection and distance measurements (**Supplementary Figure S1A-B**) that cilia always located at the cell body of proliferating cells and of neurons with differentiating neurites. Using microscopy image stacks and 3-D rendering along the full z-axis encompassing entire LUHMES cells or neurons we found the cilium to always locate in the middle sections of the stacks (**Supplementary Figure S1C; Supplementary Movie S1**). Primary cilia in LUHMES are thus ideally located for, e.g., sensory tasks concerning neighboring cells or the culture medium. The length of LUHMES cilia can reach up to 4-5 µm (our own unpublished observations). On average LUHMES cilia are thus a bit shorter than cilia from, e.g., the human retinal pigment epithelial 1 (RPE1) cell line (Pitaval et al., 2010).

### LUHMES gene expression dynamics: from the progenitor-type cell state to differentiated neurons

To monitor gene expression changes during the differentiation process, from proliferating cells to differentiated neurons, we applied the STRT RNA-sequencing method, which captures the 5’-ends of poly-A transcripts (transcript far 5’ ends, TFEs) (Islam et al., 2011, 2014; Töhönen et al., 2015; Krjutškov et al., 2016). We sampled total RNA from d0 to d6 in two independent time course experiments with 5-6 replicates per time point; in total 81 samples were sequenced (**Supplementary Table S1; Supplementary Figure S2**). After having removed 8 samples that failed quality checks and 3 technical replicates, a subtotal of 70 samples were subjected to principal component analyses (PCA) based on the expression levels of TFEs. We corrected biases between the 3 sample libraries with ComBat (Johnson et al., 2007) (**Supplementary Figure S3**). In the resulting PCA plot we confirmed that all the samples clustered based on the time points during the differentiation time course (**Figure 6A**). Notably, d0 samples (proliferating LUHMES cells) cluster separately from all other time points, indicating that gene expression changed drastically between d0 and d1, reflecting the induction of neuronal differentiation. d5 and d6 samples cluster very closely, indicating that the gene expression profile of neuronal differentiation became stable around d5, reflecting the emergence of fully differentiated neurons. Correlations between samples also supported this notion, showing low correlation between d0 and samples from other time points, but high correlation around and after d5 (**Figure 6B**).

**Figure 6:**
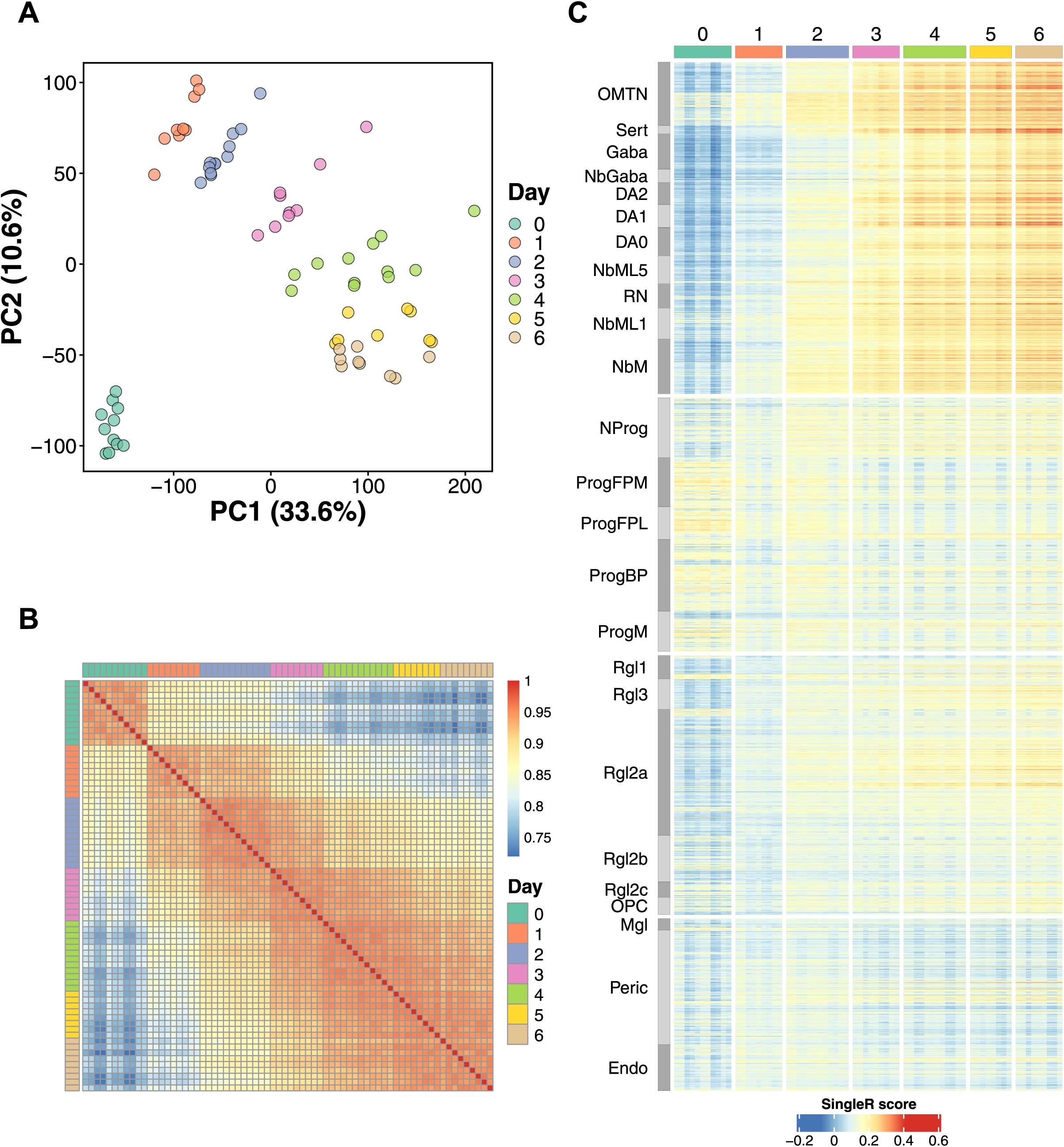
Global gene expression dynamics during the differentiation of LUHMES cells into neurons. **(A)** Principal component analysis and **(B)** Pearson correlations based on 32.483 significantly fluctuating transcript far 5’ ends (TFEs) of 70 STRT RNA-seq time course samples, representing LUHMES cells differentiating into neurons (d0 to d6). **(C)** Similarity heat map of SingleR annotation scores of different types of human fetal midbrain cells and neurons using the 70 LUHMES STRT RNA-seq time course samples as reference. Note the pronounced differences between LUHMES proliferating cells (d0) as compared to differentiating neurons (d2 to d6), and the corresponding human midbrain cells and neuron types. OMTN, oculomotor and trochlear nucleus; Sert, serotonergic neurons; Gaba, GABAergic neurons; DA, dopaminergic neurons; NbML, mediolateral neuroblasts; RN, red nucleus; NbM, medial neuroblast; NProg, neuronal progenitor; Prog, progenitor medial floor plate (FPM), lateral floor plate (FPL), basal plate (BP), midline (M); Rgl, radial glia-like cells; OPC, oligodendrocyte precursor cells; Mgl, microglia; Peric, pericytes; Endo, endothelial cells. The order of these clusters follows the order originally described in (La Manno et al., 2016).

Following previous reports about the LUHMES cell model (Delp et al., 2018; Pierce et al., 2018), we performed gene-based differential expression analysis between d0 and d6. We found 5800 significantly upregulated genes, whereas 3026 genes were downregulated (**Supplementary Figure S4**). We performed GO term enrichment analysis on these gene lists (**Supplementary Table S2**), and confirmed that terms related to ‘nervous system development’, ’vesicular transport’ or ‘axonogenesis’ were significantly enriched in the upregulated genes, appropriately reflecting that LUHMES cells differentiated into functional neurons. Interestingly, also ‘autophagy’, ’proteasome’ or ’ubiquitination’ related terms were significantly enriched in the upregulated genes. It is known that these processes play essential roles in development and cell differentiation by recycling cellular components to meet the biochemical demands (Mizushima and Levine, 2010). Autophagy and proteasome related features have also been reported for neurogenesis (Morgado et al., 2015), for neuronal differentiation (reviewed in Casares-Crespo et al., 2018), but also in connection to ciliogenesis (reviewed in Malicki and Johnson, 2017). Importantly, ‘cilium assembly’ itself was also significantly enriched in the upregulated genes. In contrast, the downregulated genes were significantly enriched with terms broadly related to ribosomal functions, in agreement with previous reports (Delp et al., 2018; Pierce et al., 2018) (**Supplementary Table S2**).

In a complementary approach we classified 5596 highly fluctuating genes into 4 clusters based on their expression pattern changes during neuronal differentiation, and identified classes of genes that were overrepresented using the core contributing genes in each cluster (shown as red lines in **Figure 7**). Genes in cluster 1 were highly expressed until d1 and monotonically downregulated after d2. Cluster 1 is highly enriched with ‘ribosomal function’ genes (**Figure 7**; **Supplementary Table S3**), indicating that translation is intensely activated on d1, when the cells begin to convert from progenitors into neurons. Genes in cluster 2 showed the highest expression on d0 and were then drastically downregulated starting on d1. Cluster 2 is highly enriched with replication and cell cycle-related genes (**Figure 7**; **Supplementary Table S3; cf. Figure 4A-D**). Genes in cluster 3 were drastically upregulated between d0 and d1. Cluster 3 is highly enriched with genes associated with transcription, transport, and catabolic processes (**Figure 7**; **Supplementary Table S3**), indicating that cells very dynamically changed their status during the early parts of neuronal differentiation. Interestingly, cluster 3 also contains sets of significantly enriched ciliary genes (**Supplementary Table S3).** Finally, we found that genes in cluster 4, most strongly upregulated from d2 to d5, were highly enriched for the terms ‘axonogenesis’ (and similar) as well as ‘cilium assembly’ (**Figure 7**; **Supplementary Table S3**). In summary, we note that highly upregulated genes during LUHMES neuronal differentiation (clusters 3 and 4) are significantly enriched with ciliary genes, including 100 ciliary genes already described in the SYSCILIA gold standard list (van Dam et al., 2013).

**Figure 7:**
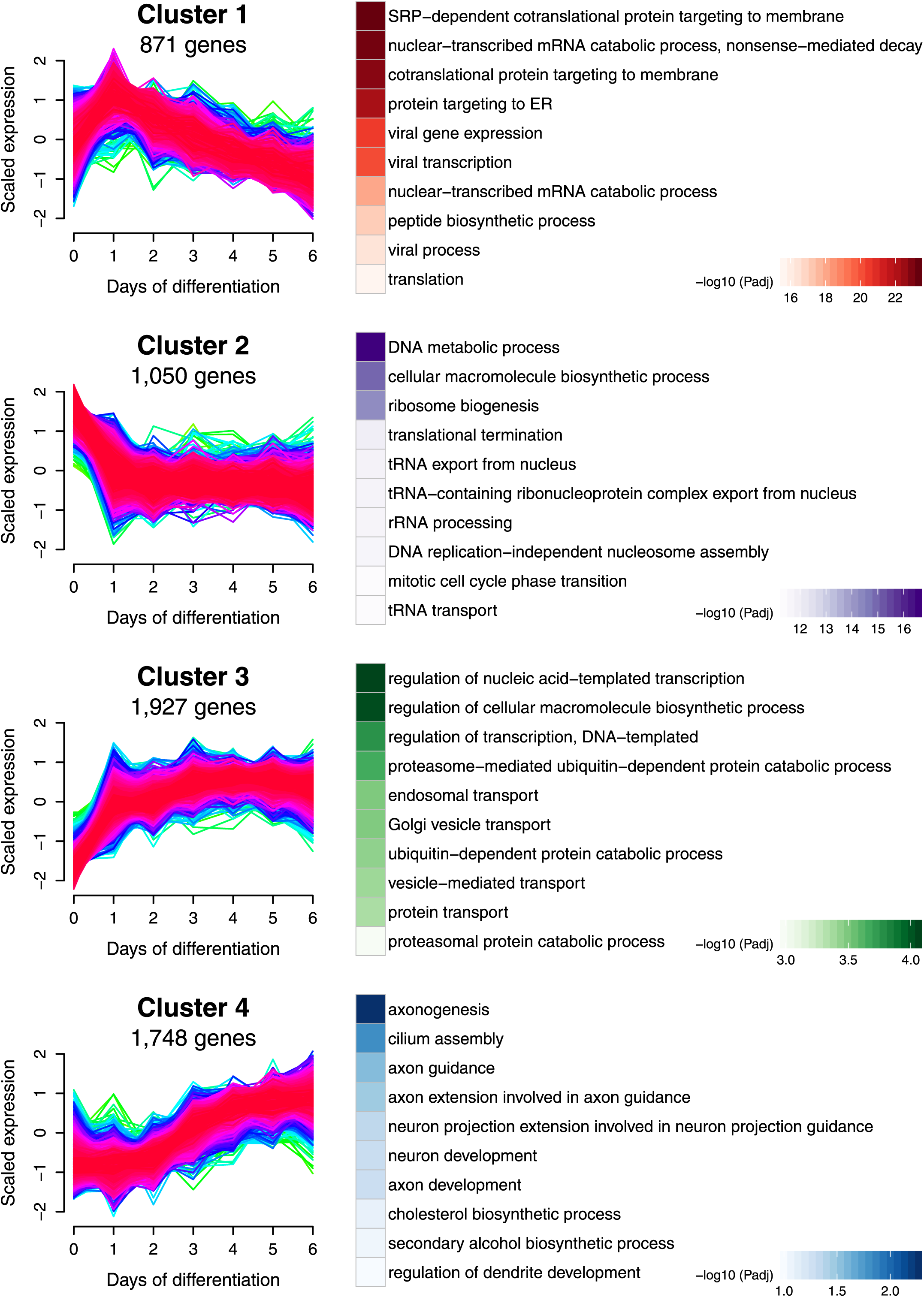
Clustering of genes by their expression patterns during the differentiation of LUHMES cells into neurons. 5596 highly fluctuating genes were clustered into four groups based on their time-course expression patterns using the ‘fuzzy c-means’ algorithm (Hathaway and Bezdek, 1986). For each cluster the heat maps on the right indicate the enrichment of GO terms of core gene sets shown as red lines in the graphs on the left. Padj, adjusted P-value.

To assess the similarity between LUHMES neuronal differentiation in culture (d0 to d6) with available in vivo data from human (neural) progenitor cells and differentiated neurons, we compared our gene expression differentiation time-course results with single cell RNA-seq data from human fetal midbrain (La Manno et al., 2016) (**Figure 6C**), using the tool SingleR, which assigns the cellular identity of single cell transcriptome data (Aran et al., 2019). We found that d0 samples (proliferating LUHMES cells) showed high similarity with progenitor-type cells from human midbrain, a similarity that was gradually lost during the neuronal differentiation time course. In contrast, LUHMES cells differentiating into neurons (d1 to d6) progressively acquired high similarity with differentiated human midbrain neurons including dopaminergic, serotonergic and GABAergic neurons. Further, LUHMES cells and neurons showed strong dissimilarity with other types of cells such as pericytes and endothelial cells (**Figure 6C**). These quantitative observations confirm that LUHMES neuronal differentiation in culture closely mimics the in vivo situation in human midbrain.

### LUHMES neuronal differentiation: RFX transcription factor binding motif activities and ciliary genes are upregulated

GO term enrichment analyses (**Supplementary Tables S2, S3**) found the term ‘cilium assembly’ (and similar) significantly enriched in the upregulated genes and in defined subsets of highly fluctuating genes (clusters 3 and 4; **Figure 7**), molecularly supporting the observed ciliogenesis during the LUHMES neuronal differentiation time course (d0 to d6). We confirmed the neuronal differentiation and ciliary gene STRT RNA-seq data with an independent method, qRT-PCR normalized against the housekeeping gene GAPDH. Pearson correlation coefficient plots demonstrated effectively identical differential gene expression patterns for a replication group of 17 genes (**Supplementary Figure S2; Supplementary Table S4**): 4 neuronal differentiation markers (DCX, MAP2, TUBA1A, TUBB3); 2 cilia and ciliopathy genes (CCDC28B, IFT20); 6 cilia and ciliopathy genes with brain or neuron phenotypes (BBS1, BBS2, CC2D2A, IFT81, PDE6D, TCTN2); 2 candidate ciliogenic transcription factor genes (RFX2, RFX5); a ciliary dyslexia candidate gene (DYX1C1/DNAAF4); and 2 neuronal Parkinson’s disease associated genes (MAPT, SNCA) (Tammimies et al., 2016; Reiter and Leroux, 2017; Lauter et al., 2018; Youn and Han, 2018). Ciliogenesis was thus found as a prominent aspect during LUHMES neuronal differentiation. Similar ‘ciliary’ observations were made for the differentiation from human neuroepithelial stem (NES) cells and induced pluripotent stem cells (iPSCs) into neurons (Bieder et al., 2020 in press).

Using the SYSCILIA gold standard list of ciliary genes (van Dam et al., 2013) we conducted gene set enrichment analysis (GSEA) by comparing each day of LUHMES neuronal differentiation with the undifferentiated state on d0. Enrichment scores (ES) were highest when we compared d0 with d5 (normalized ES = 2.57, P < 0.001) (**Figure 8A**; **Supplementary Figure S5**), demonstrating that ciliary genes were most significantly enriched in the upregulated genes on d5. This outcome is in accordance with the gene expression patterns of highly fluctuating genes in cluster 4, which showed strong upregulation prominently including d5 of differentiation (**Figure 7**). These results prompted us to explore which regulatory factors might control the expression of ciliary genes during LUHMES neuronal differentiation.

**Figure 8:**
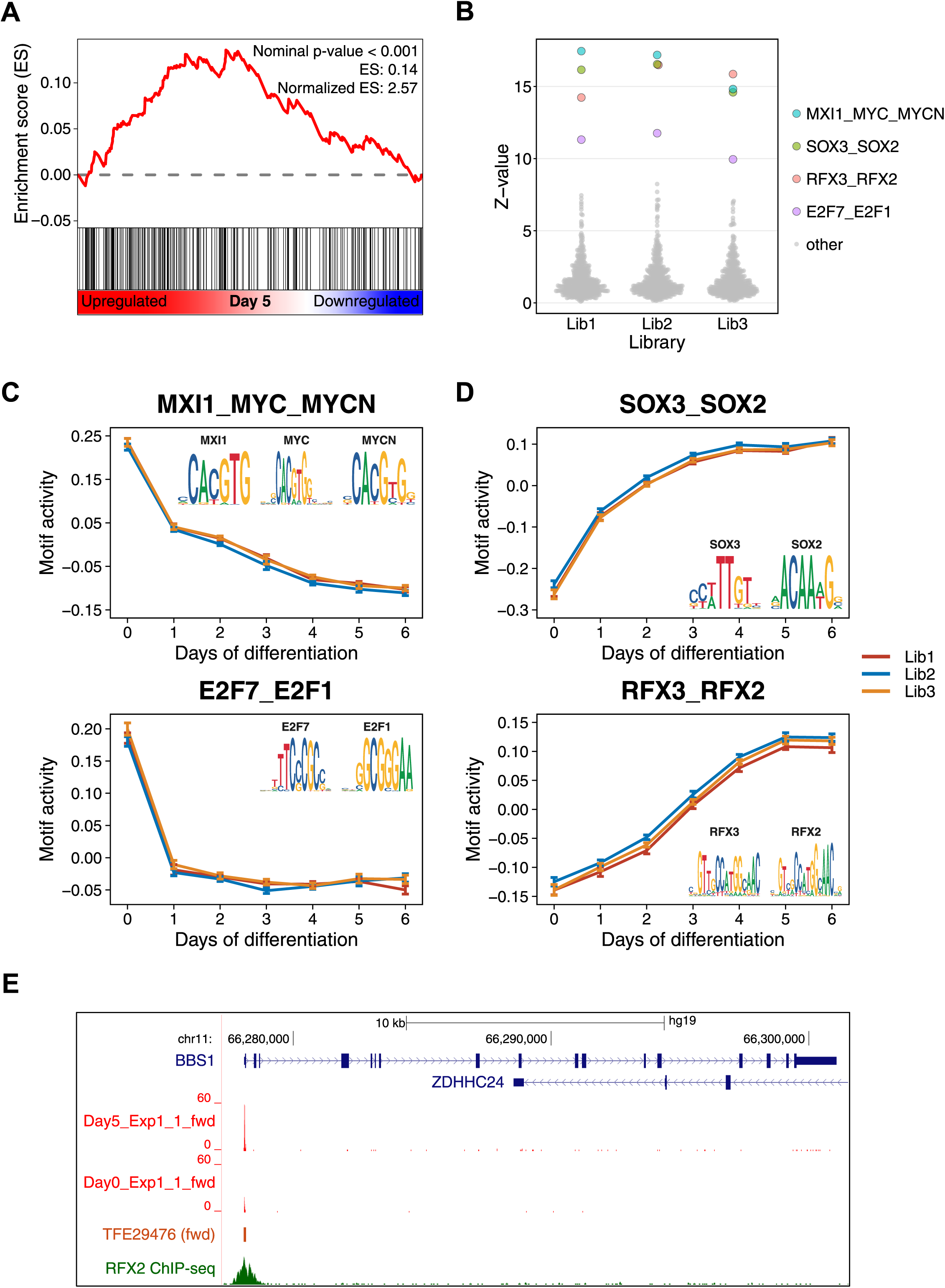
During the differentiation of LUHMES cells into neurons the activity states of ciliogenic RFX transcription factor binding motifs (X-boxes) and the expression of ciliary genes are upregulated. **(A)** Ciliary gene set enrichment analysis of differentiating LUHMES neurons (d5) compared with proliferating LUHMES cells (d0). Black vertical lines represent ranked ciliary genes based on differential gene expression comparisons between d0 and d5. See also **Supplementary Figure S5. (B)** Distribution of Z-values, which indicate TF binding motif activity changes during LUHMES differentiation into neurons. The top four motifs show remarkably high Z-values compared with all other motifs analyzed. **(C-D)** TF binding motif activity changes of MXI1_MYC_MYCN and E2F7_E2F1, and of SOX3_SOX2 and RFX3_RFX2 during LUHMES differentiation into neurons. Different line colors represent independent sequencing libraries analyzed. TF binding motifs are shown as sequence logos. **(E)** Exemplary genome browser view of the ciliary BBS1 gene region. The transcript far 5’ end TFE29476 (brown) of the BBS1 gene is significantly upregulated on d5 compared with d0 (red). TFE29476 overlaps with the RFX2 TF binding site (green). Details of all sequenced samples are listed in **Supplementary Table S1**, showing that these two samples (d0 and d5) were sequenced from the same library. fwd, forward strand.

We applied **M**otif **A**ctivity **R**esponse **A**nalysis (MARA) (FANTOM Consortium et al., 2009; Balwierz et al., 2014) to find candidate transcription factors responsible for gene expression changes during LUHMES neuronal differentiation. MARA computationally predicts key transcription factors that drive gene expression changes by integrating gene expression data with transcription factor binding motif statistics in gene promoters. MARA output consists of a binding motif activity Z-value, which quantifies the contribution of a given binding motif to the gene expression variations of its target genes across individual time points of a time course. In this analysis ‘MXI1_MYC_MYCN’, ‘SOX3_SOX2’, ‘RFX3_RFX2’, and ‘E2F7_E2F1’ showed exceptionally high Z-values in all the 3 libraries (**Figure 8B**; **Supplementary Table S5**). Whereby, the activity Z-values of ‘MXI1_MYC_MYCN’ and ‘E2F7_E2F1’ decreased drastically on d1 (**Figure 8C**). This observation is explained by the addition of tetracycline to the neuronal differentiation growth medium, which shuts down a v-myc expression cassette (cf. Materials and Methods). The E2F transcription factor family is known to play crucial roles in the control of the cell cycle, which is in accordance with the downregulation of cell cycle-related genes on d1 (cluster 2 in **Figure 7**). In contrast, activity Z-values of ‘SOX3_SOX2’ and ‘RFX3_RFX2’ increased during LUHMES neuronal differentiation (**Figure 8D**). SOX2 is known to be required for the maintenance of neural stem cells (Pevny and Nicolis, 2010), and downregulated during neuronal differentiation (Zhang and Cui, 2014). Here we also found that SOX2 expression was downregulated during LUHMES neuronal differentiation (**Supplementary Figure S2R**), indicating that SOX2 might act like a repressor for neuronal genes (Liu et al., 2014). The activity Z-value of ‘RFX3_RFX2’ strongly increased during LUHMES neuronal differentiation until d5, a signature very similar to the upregulated expression patterns of many ciliary genes (**Figure 8D; Supplementary Figure S2)**. RFX transcription factors are known to be key ciliogenic regulators in many different organisms (Senti and Swoboda, 2008; Piasecki et al., 2010; Choksi et al., 2014; Sugiaman-Trapman et al., 2018). As an example, the ciliary gene BBS1, commonly involved in a ciliopathy with brain phenotypes – Bardet-Biedl Syndrome (Mykytyn et al., 2002), was also upregulated until d5 (**Supplementary Figure S2G**). Notably, the BBS1 gene promoter has a binding site for RFX2 (Yan et al., 2013) (**Figure 8E**). For comparison, RFX3 showed a similarly upregulated expression pattern until d5, while RFX2 was most highly expressed on d1 but downregulated after d2 (**Supplementary Figure S2R**). These results suggest that during LUHMES neuronal differentiation RFX2 might be responsible for the induction of ciliogenesis early on, while RFX3 is responsible for ciliary maintenance later on. We are aware of possible redundancies though, given that the human RFX transcription factor family consists of 8 members (Sugiaman-Trapman et al., 2018) and that RFX binding motifs (X-boxes) from the different family members are very similar to each other (**Figure 8D**).

We independently confirmed the above findings by motif enrichment analysis of the differentially expressed TFEs during LUHMES neuronal differentiation. E2F and MYC binding motifs were significantly enriched in downregulated TFEs between d0 and d1. RFX binding motifs were significantly enriched in upregulated TFEs between d3 and d4, as well as between d4 and d5 (**Supplementary Table S6**). Taken together, these observations strongly suggest that RFX transcription factors play a crucial role in the regulation of ciliary genes during LUHMES neuronal differentiation.

## Discussion

We have thoroughly characterized the LUHMES ciliated human neuronal cell line. The LUHMES neuronal cell model is well established and a large body of published work demonstrates its experimental and technical usefulness for basic neurobiology research. To an array of cell biological, molecular and biochemical, including high-throughput approaches, we add a series of experimental investigations of cilia. Below we highlight important advantages and possible limitations with a focus on cilia biology.

We have appropriated and successfully optimized standardized culture conditions (Scholz et al., 2011): work with LUHMES cells and neurons is technically straightforward, and the culture conditions used do not cause any relevant losses by cell death. LUHMES cells and their cilia can be experimented with under proliferative conditions, by maintaining progenitor-type cells going through the cell cycle. Alternatively, we have differentiated LUHMES cells into functional neurons with cilia within about one week following published protocols (Lotharius et al., 2005; Scholz et al., 2011), whereby differentiated LUHMES neurons can be kept in culture up to two weeks. Immunostainings with functional neuronal differentiation markers have been performed up to d9 (Shah et al., 2016) or d13 in our hands. Neuronal physiological parameters have been assessed using both (metabolite) uptake assays and electrophysiological recordings (Scholz et al., 2011). A particular strength of the LUHMES cell model is the fact that one can completely separate cell proliferation from neuronal differentiation within one day of changing the culture medium. Such strict experimental separation greatly facilitates comparative studies of ciliary structure and function (e.g., signal transduction) between two different biological situations, while keeping an identical genetic background. The LUHMES cell model is amenable to genetic modification. Concordant with Shah et al. (2016), lipid-based transfection methods were inefficient in our hands (< 1%), while nucleofection-based methods with improved transfection efficiency (up to around 30%) are more practical, for example for CRISPR/Cas9-based genome engineering approaches (Shah et al., 2016; and our own unpublished work). One can thus examine, within a matter of weeks, the biological consequences of human brain disorder candidate mutations in a homologous system (a human neuronal cell line), while background genetic variation present in human patient material is obviously not attainable in the LUHMES cell model.

Karyotyping of the LUHMES cell model revealed a normal, female diploid set of chromosomes (Shah et al., 2016). Even though LUHMES cells and neurons are maintained in vitro in cell culture, our immunocytochemistry and detailed transcriptomics time course work demonstrated that both stages, proliferation and differentiation, showed exceptionally high similarities to the corresponding in vivo cell types, progenitors and differentiating neurons from human midbrain (La Manno et al., 2016), the LUHMES tissue of origin (Lotharius et al., 2002, 2005). Slight adjustments in LUHMES cell culture growth media even allow for driving differentiation toward a specific neuron class, like dopaminergic neurons (Scholz et al., 2011; Luk et al., 2015). Damage to and loss of dopaminergic neurons is a hallmark of Parkinson’s disease patients (reviewed in Hegarty et al., 2013). Thus, the LUHMES cell model has commonly been used in Parkinson’s disease studies and in neurotoxicity assays (Schildknecht et al., 2009; Stepkowski et al., 2015; Pierce et al., 2018; Ganjam et al., 2019; Kranaster et al., 2020). Whether there is any relevant connection between cilia malfunction and Parkinson’s disease is entirely unexplored though. Particularly our in-depth transcriptomics time course analyses of human neuronal differentiation complement and greatly extend similar transciptomics work in the mouse (Guo et al., 2019). Using the LUHMES cell model we provide step-by-step descriptions of the molecular flux from progenitor cell proliferation, to cell polarization and then neuronal differentiation, whereby we include a first description of the ciliary contributions during these neuro-developmental events.

The length of LUHMES cilia typically reaches up to 4-5 µm. This may pose limitations on (fluorescence-based) live imaging of neuronal ciliary transport (IFT) or signal transduction, or render difficult the analysis of subtle ciliary structural defects. In all experiments we have grown LUHMES cells and neurons in vitro in two-dimensional monolayer cultures. While this approach is advantageous for microscopy, it does not entirely reflect the complex three-dimensional, in vivo situation in the human brain. Further, the LUHMES cell model consists of a single cell type, progenitor cells or differentiating neurons, respectively. Again, this does not reflect the in vivo situation in the (developing) human brain, where a mixture of different cell types is present at any given time (most relevant here: La Manno et al., 2016). While brain organoid approaches (Lehmann et al., 2019) would remedy this limitation, these would make impossible the detailed time course studies presented here. LUHMES neurons have successfully been co-cultured with astrocytes (Efremova et al., 2015) and in 3-D cultures (Smirnova et al., 2016). However, brain organoids in connection to LUHMES neurons have not been reported yet.

In conclusion, we have successfully established a human neuronal cell model, LUHMES, where ciliogenesis, cilia structure and function can easily be investigated, both in the context of (neural) progenitor cell type proliferation as well as during the entire time course of differentiation into a mature, functional neuron. Particularly, the speed and one-week time frame of the LUHMES cell model, in combination with an available experimental, neuronal and ciliary, toolbox, make for an ideal test-bed to investigate (genetic) defects associated with *neuronal* cilia. LUHMES will thus be of great value for better understanding specific brain phenotypes of human ciliopathies like Joubert Syndrome, certain neurodevelopmental disruptions as observed in dyslexia and aspects of neuropsychiatric dysfunctions like in autism-spectrum disorders, for all of which strong connections to ciliary dysfunction have been uncovered (Reiter and Leroux 2017; Lauter et al., 2018; Thomas et al., 2019).

## Materials and Methods

### Cell culture media and growth conditions

**Lu**nd **h**uman **mes**encephalic (LUHMES) cells constitute a human midbrain-derived neuronal cell model. The tissue source, the ventral midbrain A9 cluster of an 8-week old female fetus, contains many dopaminergic neurons highly relevant for the development of the substantia nigra pars compacta (SNc). LUHMES cells were originally sub-cloned from MESC2.10 cells. Both the MESC2.10 and LUHMES cell lines have been developed at the Wallenberg Neuroscience Center at Lund University in Sweden (Lotharius et al., 2002, 2005). We have obtained LUHMES cells from the ATCC (https://www.lgcstandards-atcc.org; CRL-2927). Prior to all experimentation LUHMES cells were collected and the absence of mycoplasma contamination was confirmed. We used the same culture conditions as previously described (Scholz et al., 2011) with minor modifications. LUHMES cells were grown at 37°C in a humidified incubator (5% CO_2_) and maintained in vessels (typically T75 flasks) pre-coated with poly-L-ornithine hydrobromide (Sigma P3655, 50 µg/mL) and human fibronectin (Sigma F1056, 1 µg/mL). The growth medium was DMEM/F-12 Ham (Sigma D6421-6) supplemented with L-glutamine (Sigma G7513, 2.5 mM), N-2 Supplement (Gibco 17502-048, 1X) and human basic Fibroblast Growth Factor (bFGF) (Thermo Fisher Scientific 68-8785-82, 20 ng/mL). LUHMES cells were grown to 80% confluency and passaged 1:6 using diluted trypsin-EDTA (0.25%, Thermo Fisher Scientific 25200056) with PBS at 1:1 volume. For all experiments except the routine maintenance of cells, cell culture vessels (typically 6-well plates) were pre-coated with higher concentrations of ornithine (100 µg/mL) and fibronectin (10 µg/mL) to improve cell attachment to the substrate. In the growth medium for LUHMES cell proliferation, 40 ng/mL bFGF was added instead of 20 ng/mL, to accelerate cell propagation. For differentiating LUHMES cells into post-mitotic neurons (d0 to d6), cells were pre-seeded at lower densities to reach a final cell density of 1.5 × 10^5^ cells/cm^2^ during the differentiation process. 24h after pre-seeding, on d0 of LUHMES differentiation into neurons, bFGF in the growth medium was replaced by tetracycline hydrochloride (Sigma T7660, 1 µg/mL), which shuts down a v-myc expression cassette used for cell line immortalization, as was previously described for -/- growth medium conditions (no addition of cAMP and GDNF; Scholz et al., 2011).

### Immunocytochemistry

LUHMES cells and neurons were grown on coverslips (No. 1.5 round-shaped borosilicate glass, VWR 631-0150) pre-treated with concentrated hydrogen chloride (37%), washed with dH_2_O and absolute ethanol, and then stored in 70% ethanol. Coverslips were placed in each well before the vessels were coated with ornithine and fibronectin. On the day of immunostaining, the wells with coverslips were fixed with 2% paraformaldehyde (Thermo Fisher Scientific FB002) in PBS for 15 minutes at room temperature. After fixation, cells and neurons were permeabilized with 0.2% TritonX-100 in PBS for 12 minutes. The coverslips were then transferred to a humid chamber and incubated with the blocking buffer (2% bovine serum albumin, 0.1% Tween20 in PBS) for 30 minutes, followed by an overnight primary antibody incubation at +4°C and a 60 minutes secondary antibody incubation at room temperature. Cell nuclei were stained with DAPI solution (Hoechst, Invitrogen H3570, 50 µg/mL in PBS) for a few seconds and the plasma membrane with Cell Mask (Invitrogen HCS Deep Red Stain H32721, 2 ng/µL in PBS) for 30 minutes at room temperature, before the coverslips were mounted on a microscope slide with Mowiol mounting medium.

The following primary antibodies were used. Our main ciliary marker was ARL13B (rabbit, Proteintech 1711-1-AP, 1:5000). Other markers include PCNT (rabbit, Human Protein Atlas HPA019887, 1:1000), ACTUB (mouse, Sigma T7451, 1:5000) and IFT88 (rabbit, ProteinTech 13967-1-AP, 1:500). Neuronal differentiation markers were TUBB3 (Tuj1) (mouse, Covance, MMS-435P, 1:3000), PSD95 (DLG4) (rabbit, Cell Signaling 2507, 1:100), KIF17 (rabbit, Abcam ab11261, 1:2000), TH (rabbit, RnD Systems 779427, 1:50). Secondary antibodies (alexa fluor dyes) were obtained from Thermo Fisher Scientific and used at a 1:600 dilution. All primary and secondary antibodies were diluted in blocking buffer during the incubation.

### Microscopy

All images of samples used for immunocytochemistry were acquired at the Live Cell Imaging facility of the Karolinska Institute, Department of Biosciences and Nutrition (https://ki.se/en/bionut/welcome-to-the-lci-facility). A Nikon Ti-E inverted point scanning confocal microscope A1+ (Nikon Instruments) was used with a Plan Apo λ 20x NA 0.75 dry objective and a Plan Apo λ 60x NA 1.4 oil objective for higher magnifications. Image analysis was carried out using ImageJ software (Schneider et al., 2012).

### Cell densities, cell viability assays, counting of cell divisions and mitotic figures

Cell densities of cultures in six-well-plates were used to determine changes in population growth. Several different defined positions within a well were imaged on a bright field microscope (Carl Zeiss, Axiovert 200M) on consecutive days of neuronal differentiation. The number of cells and neurons counted in these defined areas was extrapolated to the entire well area to derive the cell density of one well at a given time point during the differentiation time course. Three independent quantifications were conducted.

Bright field pictures of cells and neurons were taken on a microscope equipped with epifluorescence (Carl Zeiss, Axiovert 200M) and dead cells were counted manually. Using an independent method, the same cell and neuron samples were stained by adding 2 drops/mL of NucGreen Dead 488 Ready Probes reagent (Thermo Fischer Scientific R37109), a fluorescent dye that stains cells lacking full plasma membrane integrity. After 10 minutes of incubation, fluorescent dead cells were imaged by microscopy and manually counted. Mitotic nuclei were counted on images of fixed DAPI-stained LUHMES cultures from d0 until d3 of neuronal differentiation. Immunofluorescence antibody detection (acetylated tubulin, ACTUB) was included as a differentiation marker. At least 150 nuclei were counted for each individual time point, whereby mitotic nuclei were visually identified.

### Localization of cilia and 3-D rendering

To determine the exact position of cilia on LUHMES cells and neurons, these were stained and imaged as described above using ARL13B as ciliary marker, DAPI for nuclei and Cell Mask to identify the cell body boundaries. To discriminate the Cell Mask fluorescence signal between the cell body and developing neurites, a circle was drawn around the cell nucleus (**Supplementary Figure S1A**). This circle was then considered as the edge of the cell body for subsequent quantifications. The cilium-nucleus distance (d2, d3, d4) and the cell body edge (d0, d2, d3, d4) were measured using the ImageJ line tool, starting from the center point of the nucleus as visualized by DAPI staining. For all time points a minimum of 50 ciliated cells or neurons were counted. For 3-D rendering snapshots of a series of 3-D z-axis stacks were complied using the 3D Viewer ImageJ plugin.

### Transmission electron microscopy (TEM)

TEM work was carried out at the electron microscopy facility of the Karolinska University Hospital Huddinge (https://ki.se/en/research/the-electron-microscopy-unit-emil). LUHMES cells were differentiated into neurons for three days and subsequently harvested in a cell pellet. Differentiating neurons were fixed in 2.5 % glutaraldehyde in 0.1M phosphate buffer, pH 7.4, and stored at 4°C until further use. Following fixation the cells were rinsed in 0.1M phosphate buffer prior to post-fixation in 2% osmium tetroxide in 0.1M phosphate buffer, pH 7.4, at 4°C for 2 hours. The cells were then stepwise dehydrated in ethanol followed by acetone and embedded in LX-112. Ultrathin sections (between 50-60 nm) were cut using a Leica EM UC 7 microtome and contrasted with uranyl acetate followed by lead citrate. The sections were examined in a Tecnai 10 transmission electron microscope at 100 kV. Digital images were acquired using a Veleta CCD camera.

### Drug treatments

LUHMES cells were cultured as described above and pre-seeded in 12-well plates in proliferation growth medium for 24h or overnight. At d0, the proliferation medium was changed to differentiation medium and LUHMES cells were incubated for an additional 48h (until d2 of neuronal differentiation). On d2, a SHH pathway activator, Smoothened Agonist (SAG, Merck 566660), or a SHH pathway antagonist, Cyclopamine hydrate (Sigma C4116), was added to the wells for 24h of treatment, prior to the extraction of total RNA. Both SAG and Cyclopamine (Stanton et al., 2010) were dissolved in DMSO and used at final concentrations of 500 nM and 10 µM, respectively. Control samples consisted of LUHMES cells and neurons cultured in differentiation medium for the same lengths of time, whereby only DMSO at 0.25% was added to the wells for 24h of treatment.

### Extraction of total RNA

Total RNA was isolated from LUHMES cells and neurons using a QIAGEN RNeasy Mini Kit and DNase Set, following the RNeasy Mini Handbook instructions. Initial RNA concentrations were measured with a Nanodrop ND-100 (Thermo Fischer Scientific). RNA quality was checked for an RNA integrity number > 8 (Agilent Tech 2200 Tape Station, KI Bioinformatics and Expression Analysis core facility; https://ki.se/en/bionut/bea-core-facility). Prior to library preparation for transcriptomics analyses, RNA concentrations were measured with a Qubit 3.0 Fluorometer (Thermo Fischer Scientific) and RNA samples to be used further were diluted to 10 ng/µl (+/- 0.5 ng/µL) in nuclease free-water.

### Quantitative real-time PCR (qRT-PCR)

500 ng of RNA were reverse transcribed into cDNA using a RevertAid H Minus First Strand cDNA Synthesis Kit (Thermo Fisher Scientific) with random hexamer primers, according to the manufacturer protocol. cDNA was diluted 1:10 and 2µl of diluted cDNA was used per reaction. The mRNA expression was analyzed by qRT-PCR using a 7500 Fast Real-Time PCR instrument (Applied Biosystems) and FastStart Universal SYBR Green Master reagents (4913850001, Sigma-Aldrich). All primer sequences are listed in **Supplementary Table S4**. Standard qRT-PCR runs consisted of a first passage at 95°C (10 min), followed by 40 cycles at 95°C (15 sec) and 60°C (1 min). Melting curve analyses were performed to assess whether only single, specific PCR products had been obtained. The relative fold-increase of gene expression was calculated – using the 2^-ΔΔCt method – by subtracting the Ct values of the gene of interest from the housekeeping gene GAPDH (ΔCt), and then normalizing the resulting ΔCt values against the control samples at d0 (Livak and Schmittgen, 2001).

### STRT RNA-seq library preparation and sequencing

We used 20 ng of RNA to generate three 48-plex RNA-seq libraries employing a modified STRT method with unique molecular identifiers (UMIs) (Islam et al., 2011, 2014). Briefly, RNA samples were placed in a 48-well plate in which to each well a universal primer, template-switching oligo-nucleotides, and a well-specific 6-bp barcode sequence (for sample identification) were added (Krjutškov et al., 2016). The synthesized cDNAs from these samples were then pooled into one library and amplified by single-primer PCR using a universal primer sequence. STRT library sequencing was carried out with an Illumina NextSeq 500 System, High Output (75 cycles), at the Biomedicum Functional Genomics Unit (FuGU), University of Helsinki, Finland (https://www.helsinki.fi/en/infrastructures/genome-analysis/infrastructures/biomedicum-functional-genomics-unit).

### STRT RNA-seq data preprocessing

The raw base call (BCL) files were demultiplexed with Picard (v2.10.10; http://broadinstitute.github.io/picard/) ExtractIlluminaBarcodes and IlluminaBasecallsToSam to create unaligned BAM files. These BAM files were converted to FASTQ files with Picard SamToFastq, and aligned to the human reference genome hg19, human ribosomal DNA unit (GenBank: U13369), and ERCC spike-ins (SRM 2374) with the GENCODE (v28) transcript annotation by HISAT2 (v2.1.0) (Kim et al., 2019) using the option “--dta”. Aligned BAM files were then merged with the original unaligned BAM files to create UMI-annotated BAM files by Picard MergeBamAlignment. Subsequently, these BAM files corresponding to each sample derived from 4 lanes were merged using Picard MergeSamFiles. Finally, potential PCR duplicates were marked with Picard MarkDuplicates. The resulting BAM files were processed with featureCounts (v1.6.2) (Liao et al., 2014) to assign the reads to 5’ ends of genes or TFEs with options “-s 1 --largestOverlap --ignoreDup --primary”. For gene-based analysis, uniquely mapped reads within the 5′ UTR or 500 bp upstream of the NCBI RefSeq protein-coding genes and the first 50 bp of spike-in sequences were counted. For TFE-based analysis, the mapped reads were assembled by StringTie (v1.3.3) (Pertea et al., 2015) with options “--fr -m 74 -c 10”, and those mapped reads within the first exons of the assembled transcripts were counted as previously described (Töhönen et al., 2015). FASTQ files after exclusion of duplicated reads were deposited in the ArrayExpress database at EMBL-EBI (https://www.ebi.ac.uk/arrayexpress) under the accession number E-MTAB-8908. The numbers of total and of mapped reads for each sample are summarized in **Supplementary Table S1**.

### Downstream expression analysis

Among the 81 samples from LUHMES cells and neurons evenly distributed over 3 separate libraries, 8 samples were excluded as outliers due to extremely high or low numbers of mapped reads. The bias in read counts between these 3 libraries was corrected by the ComBat function (Johnson et al., 2007) of the R (v3.6.0) package sva (v3.32.1) (Leek et al., 2012), using the remaining 73 LUHMES samples and 59 reference samples, also evenly distributed over the same 3 separate libraries, all of which were processed and sequenced in the exact same way. After removing 3 technical replicates, 70 LUHMES samples were used for follow-up analyses. Corrected read counts were normalized by the sum of spike-in reads for each sample (Katayama et al., 2013). The significance of variation (fluctuation) of normalized expression levels was evaluated by comparing them with the technical variation of spike-ins, as described in Supplementary Text S1 of Krjutškov et al. (2016). In the line plots, normalized expression values were divided by the minimum value but non-zero (5.78e-06) for visualization.

Pearson correlation coefficients between STRT RNA-seq and qRT-PCR data (**Supplementary Figure S2**) were calculated using the log2 fold change values from day 0 for each method (normalized read counts for STRT RNA-seq and 2^-ΔΔCt for qRT-PCR) per independent experiment.

Principal component analysis (PCA) was performed on significantly fluctuating TFEs (FDR-adjusted P < 0.05) across all 70 LUHMES samples, using the pca function of the R package mixOmics (v6.8.0) (Rohart et al., 2017). Pearson correlation coefficients between samples were calculated using the log2-scaled normalized expression values of significantly fluctuating TFEs. Differential expression analysis was carried out using the R package SAMstrt (v0.99.0) (Katayama et al., 2013), where we defined differentially expressed TFEs as those showing a differential expression BH-adjusted P < 0.05 with fluctuation FDR-adjusted P < 0.05. Motif enrichment on differentially expressed TFEs was analyzed using the command findMotifsGenome.pl from HOMER (v4.10.3) (Heinz et al., 2010) with the option “-size -300,100 -mask”, using all the detected TFEs as background. Clustering analysis by time-course expression pattern was carried out using the R package Mfuzz (v2.44.0) (Kumar and Futschik, 2007). A total of 5596 genes that showed highly significant fluctuation (FDR-adjusted P < 1e-10) for each experiment and high correlation between two experiments (Pearson correlation > 0.7) were classified into 4 clusters, which were determined based on the minimum centroid distance. Gene ontology (GO) term enrichment analysis was performed using the R package enrichR (v2.1) (Kuleshov et al., 2016). Core genes in each cluster for the enrichment analysis were extracted based on high membership scores (> 0.75) using the acore function of the Mfuzz package. Gene set enrichment analysis (GSEA) was performed with GSEA (v3.0; http://software.broadinstitute.org/gsea/) using the GSEAPreranked tool (Subramanian et al., 2005). Here, genes were ranked based on the combination of differential expression score and fold changes of their expression levels, and were then compared with the gene set of 302 ciliary genes from the SYSCILIA gold standard (SCGSv1) (van Dam et al., 2013). GSEA plots were made using ReplotGSEA.R (https://github.com/PeeperLab/Rtoolbox/blob/master/R/ReplotGSEA.R) with minor modifications. Integrative analysis with fetal midbrain single cell RNA-seq data (GSE76381) (La Manno et al., 2016) was carried out with the CreateSinglerSeuratObject function of the R package SingleR (v1.0.1) (Aran et al., 2019) and Seurat (v3.0.2) (Satija et al., 2015), using the gene-based normalized expression data of LUHMES samples as the reference data set.

After removing duplicated, non-primary, and unmapped reads from the BAM files followed by sorting with SAMtools (v1.9), these processed BAM files from each library were subjected to the ISMARA (The Integrated System for Motif Activity Response Analysis) online tool (https://ismara.unibas.ch/mara/) (Balwierz et al., 2014) for the motif activity response analysis. Sequence logos were generated using the R package ggseqlogo (v0.1) (Wagih, 2017). All the analyses scripts are available upon request.

## Acknowledgements

Parts of this study were performed at the Karolinska Institute Live Cell Imaging facility supported by grants from the Knut and Alice Wallenberg Foundation, the Swedish Research Council, the Centre for Innovative Medicine and the Jonasson Center at the Swedish Royal Institute of Technology. We acknowledge technical and personnel support from the electron microscopy facility of the Karolinska University Hospital Huddinge. We acknowledge BEA, the Bioinformatics and Expression Analysis core facility, which is supported by the board of research at the Karolinska Institute and the research committee at the Karolinska Hospital. The Biomedicum Functional Genomics Unit at the University of Helsinki provided sequencing services. All computation for this work was performed on resources provided through the Uppsala Multidisciplinary Center for Advanced Computational Science (UPPMAX) under project contract SNIC 2017/7-317.

## Author Contributions

Project conceptualization and planning: GL, JK, PS

Experimentation and methodology: GL, AC, MY, DST, SE, SS

Data analysis: GL, AC, MY, DST, SK, JK, PS

Critical resources and reagents: GL, MY, SK

Writing and illustrations – draft: GL, AC, MY, PS

Writing and illustrations – editing and review: GL, AC, MY, JK, PS

Supervision and project management: JK, PS

Funding acquisition: JK, PS

## Data Availability

See Materials and Methods for this part.

## Supplementary Information

Supplementary information is available online at (Insert web link here).

## Supplementary Materials

**Supplementary Figure S1:**
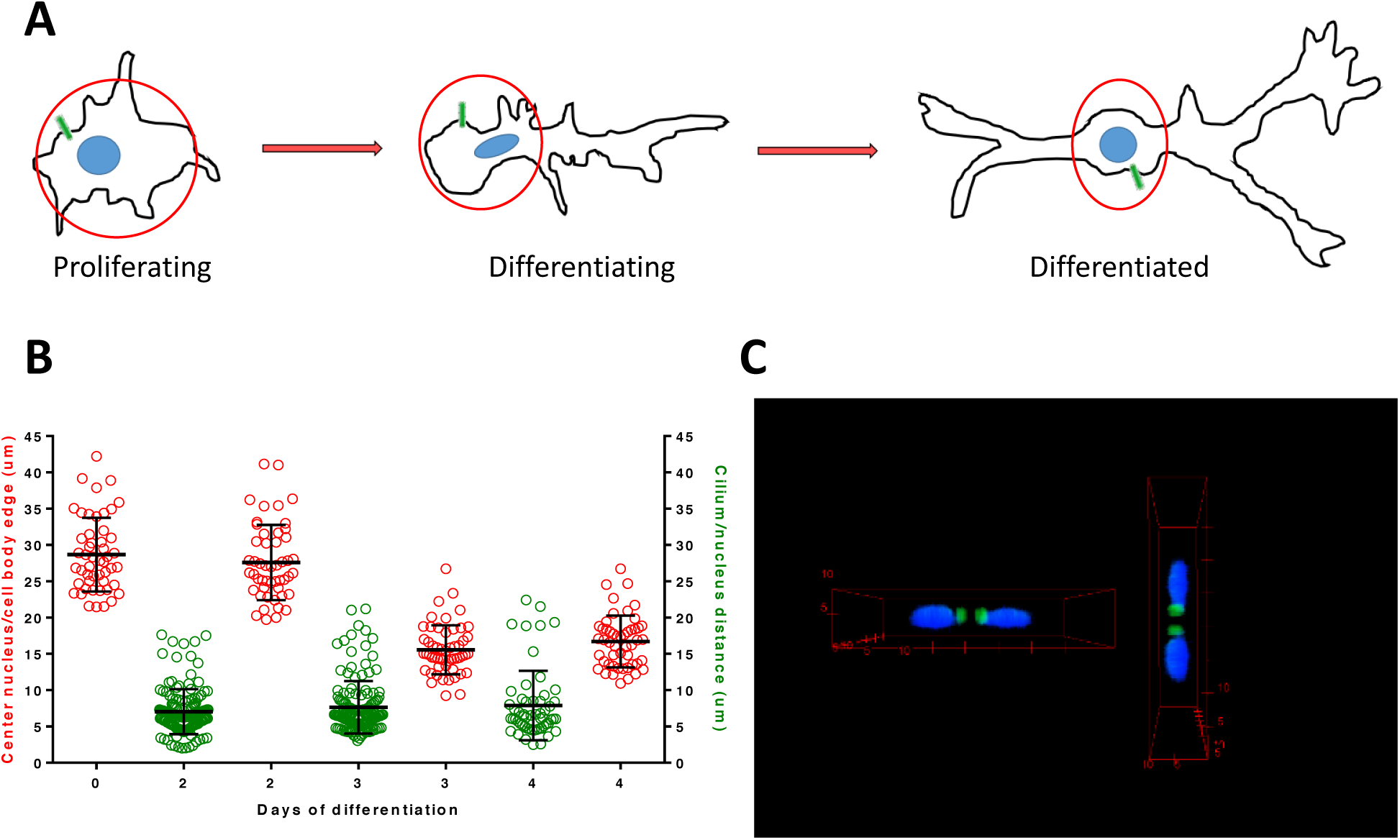
Location of cilia in differentiating LUHMES neurons. **(A)** Schematic representation of LUHMES cells and neurons at proliferating (d0), differentiating (d3) and differentiated (d6) stages. Primary cilia are depicted in green and nuclei in blue, whereby the cell body is contained within the red circle. **(B)** The distances from the center of the nucleus to the boundaries of the cell body (transition-zone-to-neurites) at d0, d2, d3, d4 (in red) are compared to the distances from the center of the nucleus to the cilium at d2, d3, d4 (in green). Cilia in differentiating LUHMES neurons are located at the cell body. **(C)** A 3-D rendering of cilia location is provided along the z-axis of stacked images. The same rendering is shown from two different viewpoints. Cilia, stained with ARL13B (green), typically appear in the middle sections of a z-stack. Nuclei are counterstained with DAPI (blue). See also **Supplementary Movie S1**. All distances in **(B-C)** are in µm.

**Supplementary Figure S2.**
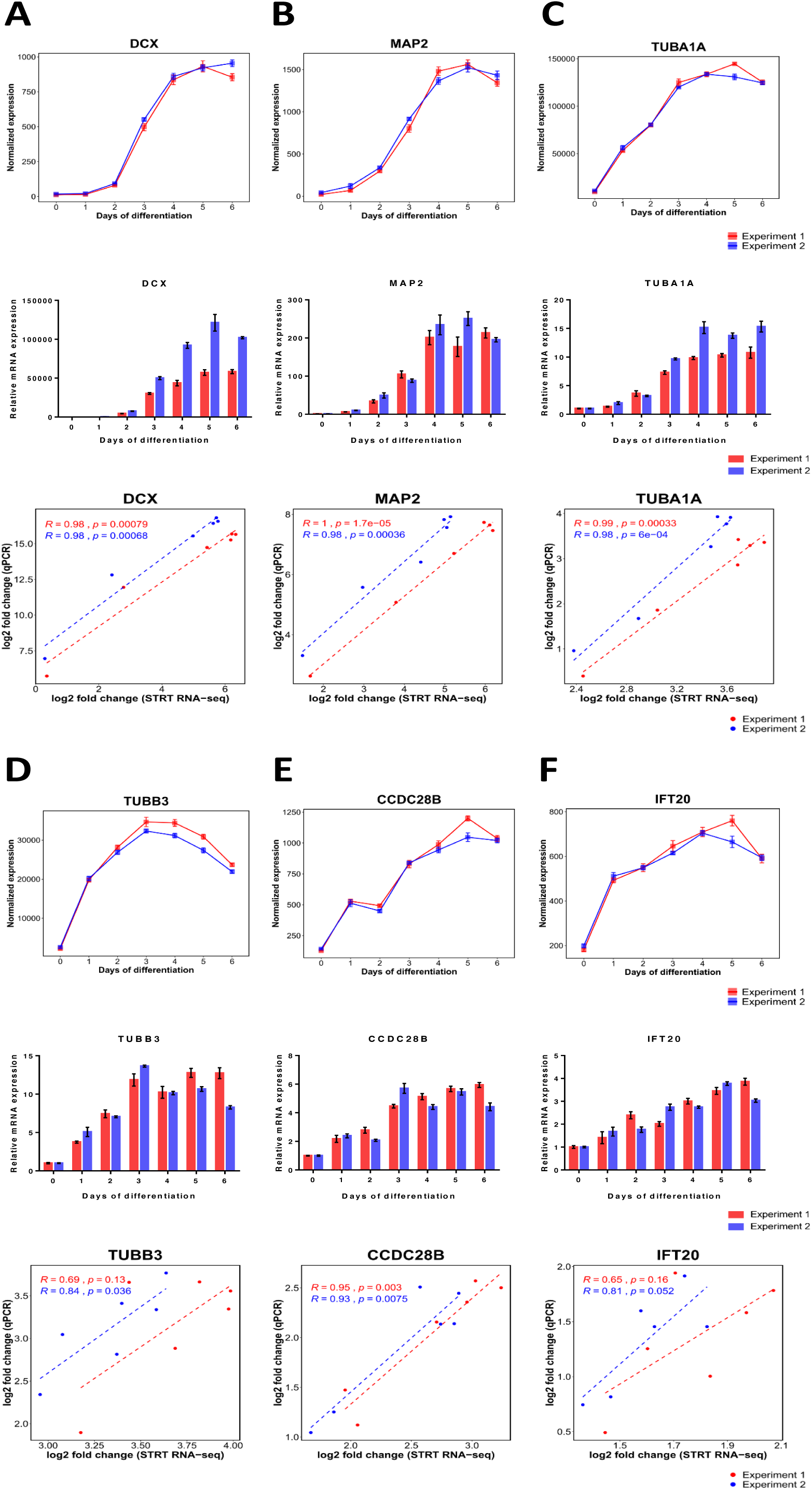
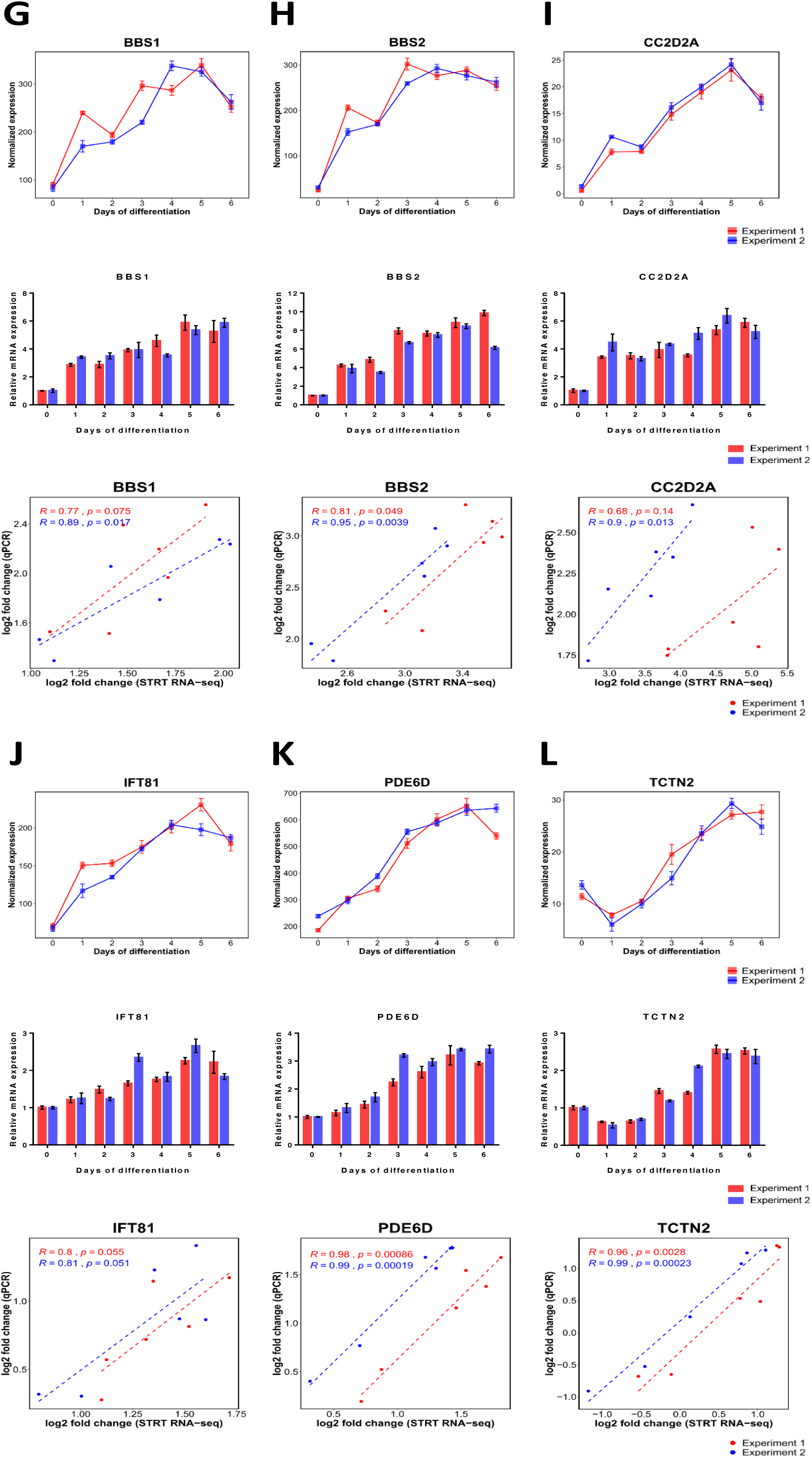
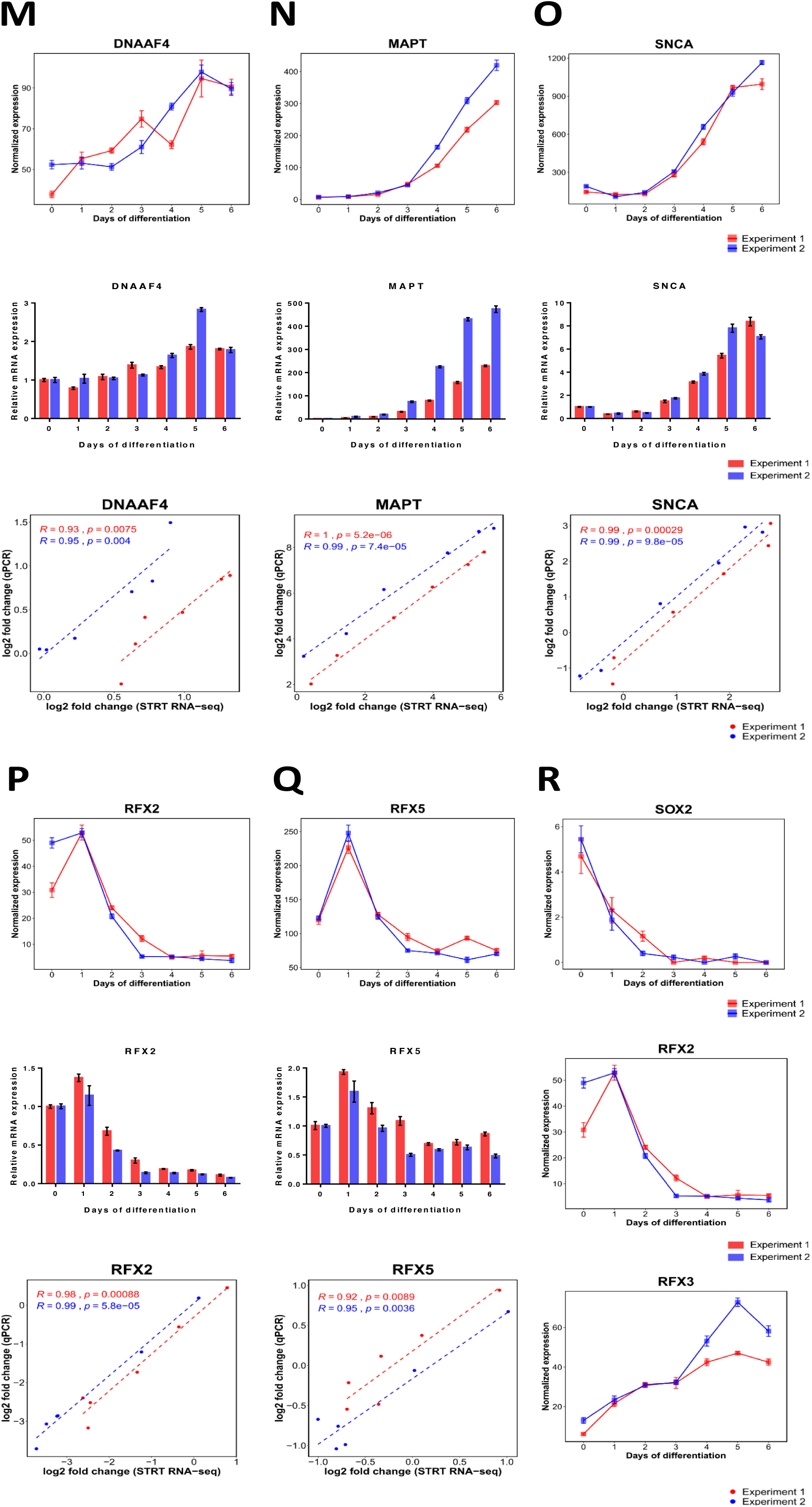
Gene expression profiles of LUHMES cells differentiating into neurons. Gene expression was assessed by STRT RNA-seq from d0 to d6 and validated by quantitative real-time PCR (qRT-PCR). Gene expression data using both methods were correlated as log_2_ fold change in STRT RNA-seq versus log_2_ fold change in qRT-PCR. R refers to the Pearson correlation coefficient. Data were normalized to the housekeeping gene GAPDH and displayed as the mean +/- SEM from two independent experiments (red and blue). **(A-Q)** Upper panels show STRT RNA-seq time course data; middle panels show qRT-PCR time course data; lower panels show the correlation plots, respectively. Differential gene expression patterns of **(A-D)** neuronal marker genes DCX, MAP2, TUBA1A, TUBB3; **(E-F)** ciliary and ciliopathy genes CCDC28B, IFT20; **(G-L)** ciliary and ciliopathy genes with associated brain phenotypes BBS1, BBS2, CC2D2A, IFT81, PDE6D, TCTN2; **(M)** the dyslexia candidate gene DNAAF4 (DYX1C1); **(N-O)** Parkinson’s disease candidate genes MAPT, SNCA; **(P-Q)** ciliogenic transcription factor genes RFX2, RFX5. **(R)** STRT RNA-seq gene expression levels of the transcription factor genes SOX2, RFX2, and RFX3 during LUHMES cells differentiating into neurons.

**Supplementary Figure S3:**
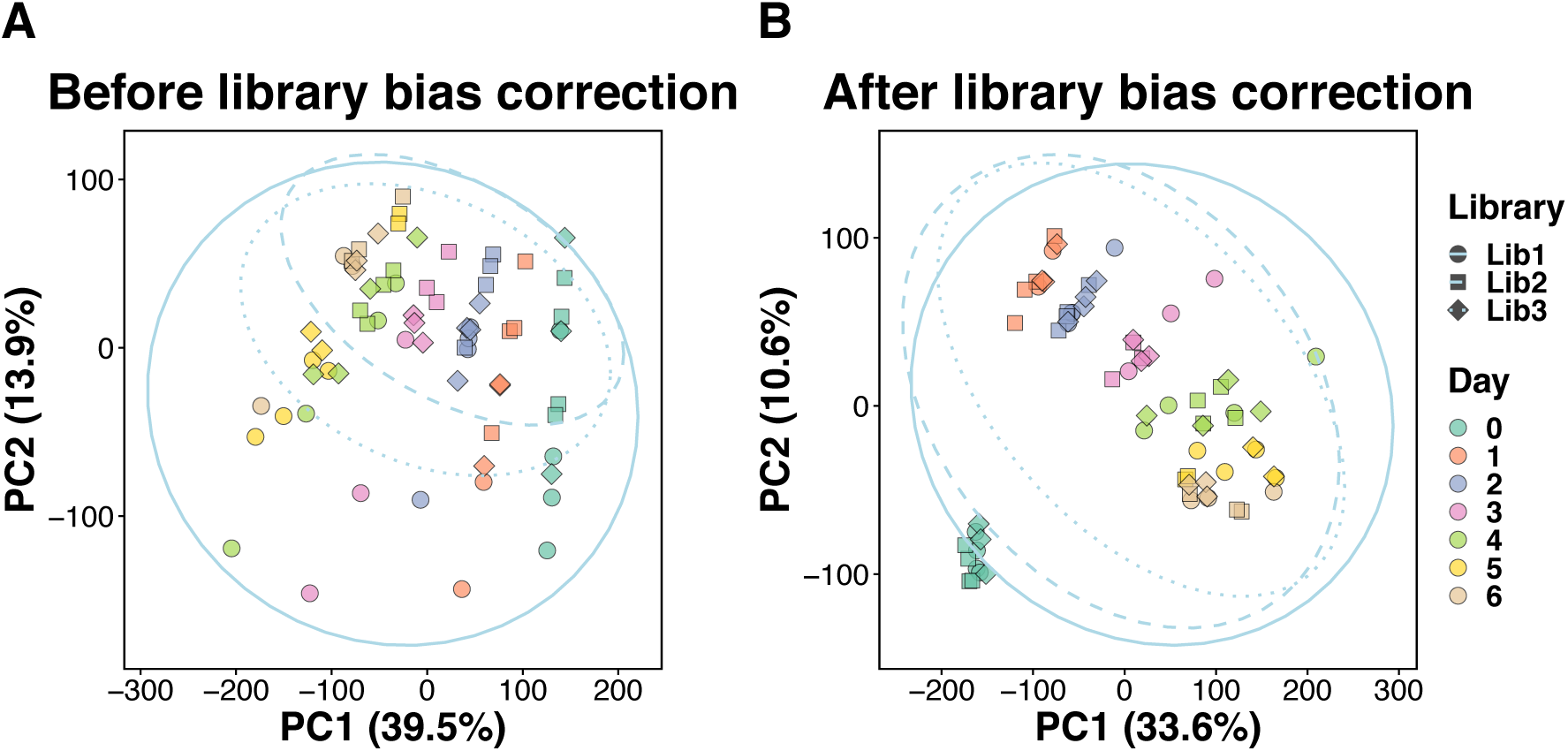
Principal component analysis (PCA) of LUHMES differentiation time course samples before and after library bias correction. PCA plots covering 32.483 significantly fluctuating transcript far 5’ ends (TFEs) of 70 STRT RNA-seq time course samples, representing LUHMES cells differentiating into neurons (d0 to d6), before **(A)** and after **(B)** library bias correction. Solid, dashed, and dotted lines represent the 95% confidence interval ellipses for each of the three independent libraries used.

**Supplementary Figure S4:**
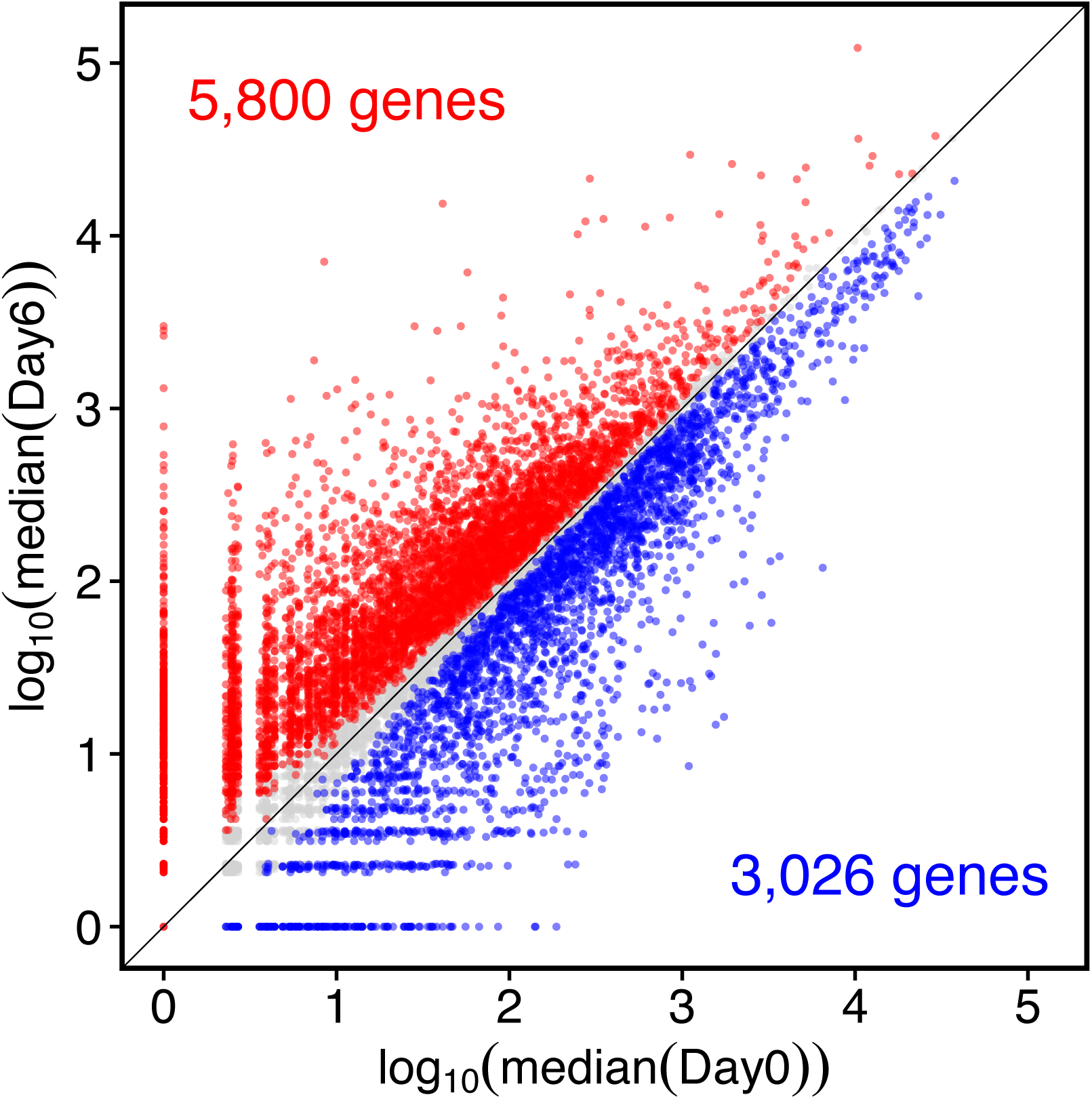
Scatter plot of STRT RNA-seq gene expression levels from LUHMES cells differentiating into neurons (d0 versus d6). Red dots represent significantly upregulated genes, while blue dots represent significantly downregulated genes on d6 (differentiated LUHMES neurons) versus d0 (proliferating LUHMES cells).

**Supplementary Figure S5:**
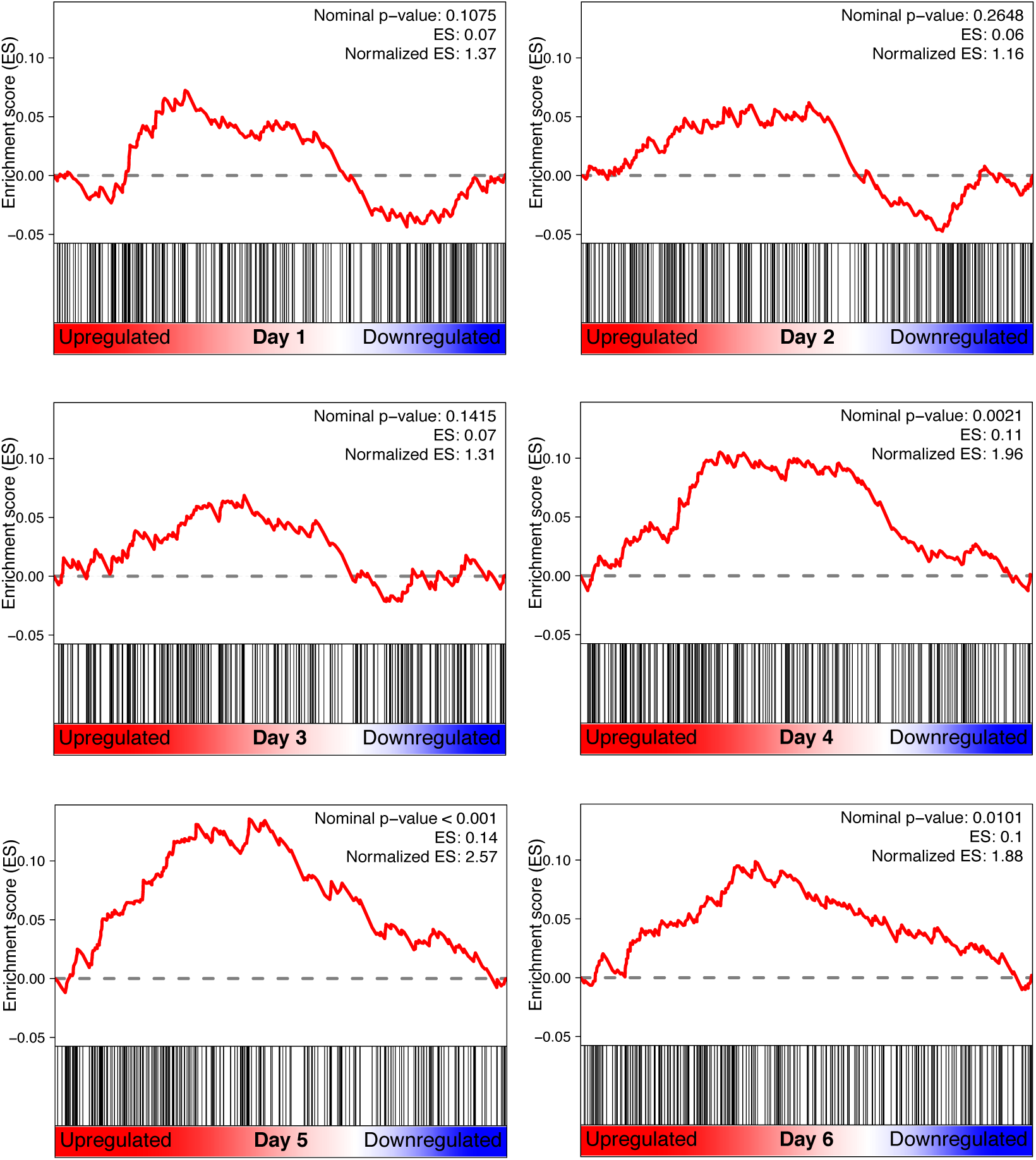
Ciliary gene set enrichment analysis (GSEA) of differentiating LUHMES neurons (d1-d6) compared with proliferating LUHMES cells (d0). Black vertical lines represent ranked ciliary genes based on differential gene expression comparisons between d0 and the respective day indicated in each graph. Enrichment scores (ES) are highest on d5 of differentiation. See also **Figure 8A**.

***Supplementary Tables*** (All are provided as separate Excel files)

**Supplementary Table S1:** Summary of STRT RNA-seq sequencing output.

**Supplementary Table S2:** Gene ontology (GO) enrichment analysis of significantly up- and downregulated genes between day 0 and day 6 of the LUHMES neuronal differentiation time course.

**Supplementary Table S3:** Gene ontology (GO) enrichment analysis of the core genes in the four clusters that are based on LUHMES neuronal differentiation time course expression patterns.

**Supplementary Table S4:** Differentially regulated genes during LUHMES neuronal differentiation (d0-d6) as determined by both STRT RNA-seq and qRT-PCR, including functional assignments, descriptions and the list of SYBR Green primers used for qRT-PCR; with the exception of the genes GLI1, HHIP and PTCH1, whose expression was quantified by qRT-PCR to assess their involvement in the SHH signaling pathway.

**Supplementary Table S5:** List of Z-values, which indicate transcription factor binding motif activity changes during LUHMES differentiation into neurons. Motif hits are listed separately for each of the three independent libraries used.

**Supplementary Table S6:** Transcription factor binding motif enrichment analysis of differentially expressed transcript far 5’ ends (TFEs) between neighboring days of LUHMES differentiation into neurons. Only the top five significantly enriched motifs are listed.

***Supplementary Movie*** (This movie is provided as separate files in both gif and mp4 format)

**Supplementary Movie S1** is a complement to **Supplementary Figure S1** – see there for details.

## References

Anvarian, Z., Mykytyn, K., Mukhopadhyay, S., Pedersen, L. B. and Christensen, S. T. (2019). Cellular signalling by primary cilia in development, organ function and disease. Nat. Rev. Nephrol. 15(4), 199–219. doi:10.1038/s41581-019-0116-9.

Aran, D., Looney, A. P., Liu, L., Wu, E., Fong, V., Hsu, A., Chak, S., Naikawadi, R. P., Wolters, P. J., Abate, A. R., et al. (2019). Reference-based analysis of lung single-cell sequencing reveals a transitional profibrotic macrophage. Nat. Immunol. 20(2), 163–172. doi:10.1038/s41590-018-0276-y.

Arellano, J. I., Guadiana, S. M., Breunig, J. J., Rakic, P. and Sarkisian, M. R. (2012). Development and distribution of neuronal cilia in mouse neocortex. J. Comp. Neurol. 520(4), 848–873. doi:10.1002/cne.22793.

Balwierz, P. J., Pachkov, M., Arnold, P., Gruber, A. J., Zavolan, M. and van Nimwegen, E. (2014). ISMARA: automated modeling of genomic signals as a democracy of regulatory motifs. Genome Res. 24(5), 869–884. doi:10.1101/gr.169508.113.

Bieder, A., Yoshihara, M., Katayama, S., Krjutškov, K., Falk, A., Kere, J. and Tapia-Páez, I. (2020). Dyslexia candidate gene and ciliary gene expression dynamics during human neuronal differentiation. Mol. Neurobiol., in press. doi:10.1007/s12035-020-01905-6.

Bishop, G. A., Berbari, N. F., Lewis, J. and Mykytyn, K. (2007). Type III adenylyl cyclase localizes to primary cilia throughout the adult mouse brain. J. Comp. Neurol. 505(5), 562–571. doi:10.1002/cne.21510.

Caspary, T., Larkins, C. E. and Anderson, K. V. (2007). The Graded Response to Sonic Hedgehog Depends on Cilia Architecture. Dev. Cell. 12(5), 767–778. doi:10.1016/j.devcel.2007.03.004.

Caspary, T., Marazziti, D. and Berbari, N. F. (2016). Methods for Visualization of Neuronal Cilia. Methods Mol. Biol. 1454, 203–214. doi:10.1007/978-1-4939-3789-9_13.

Casares-Crespo, L., Calatayud-Baselga, I., García-Corzo, L. and Mira, H. (2018). On the Role of Basal Autophagy in Adult Neural Stem Cells and Neurogenesis. Front. Cell. Neurosci. 12, 339. doi:10.3389/fncel.2018.00339.

Choksi, S. P., Lauter, G., Swoboda, P. and Roy, S. (2014). Switching on cilia: transcriptional networks regulating ciliogenesis. Development. 141(7), 1427–1441. doi:10.1242/dev.074666.

Delp, J., Gutbier, S., Cerff, M., Zasada, C., Niedenführ, S., Zhao, L., Smirnova, L., Hartung, T., Borlinghaus, H., Schreiber, F., et al. (2018). Stage-specific metabolic features of differentiating neurons: Implications for toxicant sensitivity. Toxicol. Appl. Pharmacol. 354, 64–80. doi:10.1016/j.taap.2017.12.013.

Efremova, L., Schildknecht, S., Adam, M., Pape, R., Gutbier, S., Hanf, B., Bürkle, A. and Leist, M. (2015). Prevention of the degeneration of human dopaminergic neurons in an astrocyte co-culture system allowing endogenous drug metabolism. Br. J. Pharmacol. 172(16), 4119–4132. doi:10.1111/bph.13193.

FANTOM Consortium, Suzuki, H., Forrest, A. R., van Nimwegen, E., Daub, C. O., Balwierz, P. J., Irvine, K. M., Lassmann, T., Ravasi, T., Hasegawa, Y., et al. (2009). The transcriptional network that controls growth arrest and differentiation in a human myeloid leukemia cell line. Nat. Genet. 41(5), 553–562. doi:10.1038/ng.375.

Ferent, J., Constable, S., Gigante, E. D., Yam, P. T., Mariani, L. E., Legué, E., Liem, K. F. Jr, Caspary, T. and Charron, F. (2019). The Ciliary Protein Arl13b Functions Outside of the Primary Cilium in Shh-Mediated Axon Guidance. Cell Rep. 29(11), 3356–3366. doi:10.1016/j.celrep.2019.11.015.

Ford, M. J., Yeyati, P. L., Mali, G. R., Keighren, M. A., Waddell, S. H., Mjoseng, H. K., Douglas, A. T., Hall, E. A., Sakaue-Sawano, A., Miyawaki, A., et al. (2018). A Cell/Cilia Cycle Biosensor for Single-Cell Kinetics Reveals Persistence of Cilia after G1/S Transition Is a General Property in Cells and Mice. Dev. Cell. 47(4), 509–523. doi:10.1016/j.devcel.2018.10.027.

Franker, M. A., Esteves Da Silva, M., Tas, R. P., Tortosa, E., Cao, Y., Frias, C. P., Janssen, A. F. J., Wulf, P. S., Kapitein, L. C. and Hoogenraad, C. C. (2016). Three-Step Model for Polarized Sorting of KIF17 into Dendrites. Curr. Biol. 26(13), 1705–1712. doi:10.1016/j.cub.2016.04.057.

Ganjam, G. K., Bolte, K., Matschke, L. A., Neitemeier, S., Dolga, A. M., Höllerhage, M., Höglinger, G. U., Adamczyk, A., Decher, N., Oertel, W. H., et al. (2019). Mitochondrial damage by α-synuclein causes cell death in human dopaminergic neurons. Cell Death Dis. 10(11), 865. doi:10.1038/s41419-019-2091-2.

Garcia, G., Raleigh, D. R. and Reiter, J. F. (2018). How the Ciliary Membrane Is Organized Inside-Out to Communicate Outside-In. Curr. Biol. 28(8), R421–R434. doi:10.1016/j.cub.2018.03.010.

Goetz, S. C. and Anderson, K. V. (2010). The primary cilium: a signalling centre during vertebrate development. Nat. Rev. Genet. 11(5), 331–344. doi:10.1038/nrg2774.

Guo, J., Otis, J. M., Suciu, S. K., Catalano, C., Xing, L., Constable, S., Wachten, D., Gupton, S., Lee, J., Lee, A., et al. (2019). Primary Cilia Signaling Promotes Axonal Tract Development and Is Disrupted in Joubert Syndrome-Related Disorders Models. Dev. Cell. 51(6), 759–774. doi:10.1016/j.devcel.2019.11.005.

Harischandra, D. S., Rokad, D., Ghaisas, S., Verma, S., Robertson, A., Jin, H., Anantharam, V., Kanthasamy, A. and Kanthasamy, A. G. (2020). Enhanced differentiation of human dopaminergic neuronal cell model for preclinical translational research in Parkinsons disease. Biochim. Biophys. Acta Mol. Basis Dis. 1866(4), 165533. doi:10.1016/j.bbadis.2019.165533.

Hathaway, R. J. and Bezdek, J. C. (1986). Local convergence of the fuzzy c-Means algorithms. Pattern Recognit. 19(6), 477–480. doi:10.1016/0031-3203(86)90047-6.

Hegarty, S. V., Sullivan, A. M. and O’Keeffe, G. W. (2013). Midbrain dopaminergic neurons: A review of the molecular circuitry that regulates their development. Dev. Biol. 379(2), 123–138. doi:10.1016/j.ydbio.2013.04.014.

Heinz, S., Benner, C., Spann, N., Bertolino, E., Lin, Y. C., Laslo, P., Cheng, J. X., Murre, C., Singh, H. and Glass, C. K. (2010). Simple Combinations of Lineage-Determining Transcription Factors Prime cis-Regulatory Elements Required for Macrophage and B Cell Identities. Mol. Cell. 38(4), 576–589. doi:10.1016/j.molcel.2010.05.004.

Hunt, C., Schenker, L. and Kennedy, M. (1996). PSD-95 is associated with the postsynaptic density and not with the presynaptic membrane at forebrain synapses. J. Neurosci. 16(4), 1380–1388. doi:10.1523/jneurosci.16-04-01380.1996.

Ishikawa, H. and Marshall, W. F. (2011). Ciliogenesis: building the cells antenna. Nat. Rev. Mol. Cell Biol. 12(4), 222–234. doi:10.1038/nrm3085.

Islam, S., Kjallquist, U., Moliner, A., Zajac, P., Fan, J.-B., Lönnerberg, P. and Linnarsson, S. (2011). Characterization of the single-cell transcriptional landscape by highly multiplex RNA-seq. Genome Res. 21(7), 1160–1167. doi:10.1101/gr.110882.110.

Islam, S., Zeisel, A., Joost, S., La Manno, G., Zajac, P., Kasper, M., Lönnerberg, P. and Linnarsson, S. (2014). Quantitative single-cell RNA-seq with unique molecular identifiers. Nat. Methods. 11(2), 163–166. doi:10.1038/nmeth.2772.

Johnson, W. E., Li, C. and Rabinovic, A. (2007). Adjusting batch effects in microarray expression data using empirical Bayes methods. Biostatistics. 8(1), 118–127. doi:10.1093/biostatistics/kxj037.

Joukov, V. and De Nicolo, A. (2019). The Centrosome and the Primary Cilium: The Yin and Yang of a Hybrid Organelle. Cells. 8(7), E701. doi:10.3390/cells8070701.

Katayama, S., Töhönen, V., Linnarsson, S. and Kere, J. (2013). SAMstrt: statistical test for differential expression in single-cell transcriptome with spike-in normalization. Bioinformatics. 29(22), 2943–2945. doi:10.1093/bioinformatics/btt511.

Kim, D., Paggi, J. M., Park, C., Bennett, C. and Salzberg, S. L. (2019). Graph-based genome alignment and genotyping with HISAT2 and HISAT-genotype. Nat. Biotechnol. 37(8), 907–915. doi:10.1038/s41587-019-0201-4.

Kranaster, P., Karreman, C., Dold, J. E. G. A., Krebs, A., Funke, M., Holzer, A. K., Klima, S., Nyffeler, J., Helfrich, S., Wittmann, V., et al. (2020). Time and space-resolved quantification of plasma membrane sialylation for measurements of cell function and neurotoxicity. Arch. Toxicol. 94(2), 449–467. doi:10.1007/s00204-019-02642-z.

Krjutškov, K., Katayama, S., Saare, M., Vera-Rodriguez, M., Lubenets, D., Samuel, K., Laisk-Podar, T., Teder, H., Einarsdottir, E., Salumets, A., et al. (2016). Single-cell transcriptome analysis of endometrial tissue. Human Reprod. 31(4), 844–853. doi:10.26226/morressier.573c1512d462b80296c98648.

Kuleshov, M. V., Jones, M. R., Rouillard, A. D., Fernandez, N. F., Duan, Q., Wang, Z., Koplev, S., Jenkins, S. L., Jagodnik, K. M., Lachmann, A., et al. (2016). Enrichr: a comprehensive gene set enrichment analysis web server 2016 update. Nucleic Acids Res. 44(W1), W90–W97. doi:10.1093/nar/gkw377.

Kumar, L. and Futschik, M. E. (2007). Mfuzz: A software package for soft clustering of microarray data. Bioinformation. 2(1), 5–7. doi:10.6026/97320630002005.

La Manno, G., Gyllborg, D., Codeluppi, S., Nishimura, K., Salto, C., Zeisel, A., Borm, L. E., Stott, S. R. W., Toledo, E. M., Villaescusa, J. C., et al. (2016). Molecular Diversity of Midbrain Development in Mouse, Human, and Stem Cells. Cell. 167(2), 566–580. doi:10.1016/j.cell.2016.09.027.

Lauter, G., Swoboda, P. and Tapia-Páez, I. (2018). Chapter 1: Cilia in Brain Development and Disease. Cilia: Development and Disease (ed. P. Goggolidou), pp. 1-35. CRC Press, Taylor and Francis Publishers, Boca Raton, Florida, USA. ISBN: 9781498703680.

Lee, J.-M., Adler, J. and Black, I. (1990). Regulation of neurotransmitter expression by a membrane-derived factor. Exp. Neurol. 108(2), 109–113. doi:10.1016/0014-4886(90)90016-l.

Leek, J. T., Johnson, W. E., Parker, H. S., Jaffe, A. E. and Storey, J. D. (2012). The sva package for removing batch effects and other unwanted variation in high-throughput experiments. Bioinformatics. 28(6), 882–883. doi:10.1093/bioinformatics/bts034.

Lehmann, R., Lee, C. M., Shugart, E. C., Benedetti, M., Charo, R. A., Gartner, Z., Hogan B., Knoblich, J., Nelson, C. M. and Wilson, K. M. (2019). Human organoids: a new dimension in cell biology. Mol. Biol. Cell. 30(10), 1129–1137. doi:10.1091/mbc.e19-03-0135.

Liao, Y. K., Smyth, G. K. and Shi, W. K. (2014). featureCounts: an efficient general purpose program for assigning sequence reads to genomic features. Bioinformatics. 30(7), 923–930. doi:10.1093/bioinformatics/btt656.

Liu, H., Kiselevaa, A. A. and Golemis E. A. (2018). Ciliary Signalling in Cancer. Nat. Rev. Cancer 18(8), 511–524. doi:10.1038/s41568-018-0023-6.

Liu, Y.-R., Laghari, Z. A., Novoa, C. A., Hughes, J., Webster, J. R., Goodwin, P. E., Wheatley, S. P. and Scotting, P. J. (2014). Sox2 acts as a transcriptional repressor in neural stem cells. BMC Neurosci. 15(1), 95. doi:10.1186/1471-2202-15-95.

Livak, K. J. and Schmittgen, T. D. (2001). Analysis of Relative Gene Expression Data Using Real-Time Quantitative PCR and the 2−ΔΔCT Method. Methods. 25(4), 402–408. doi:10.1006/meth.2001.1262.

Lotharius, J., Barg, S., Wiekop, P., Lundberg, C., Raymon, H. K. and Brundin, P. (2002). Effect of Mutant α-Synuclein on Dopamine Homeostasis in a New Human Mesencephalic Cell Line. J. Biol. Chem. 277(41), 38884–38894. doi:10.1074/jbc.M205518200.

Lotharius, J., Falsig, J., van Beek, J., Payne, S., Dringen, R., Brundin, P. and Leist, M. (2005). Progressive Degeneration of Human Mesencephalic Neuron-Derived Cells Triggered by Dopamine-Dependent Oxidative Stress Is Dependent on the Mixed-Lineage Kinase Pathway. J. Neurosci. 25(27), 6329–6342. doi:10.1523/jneurosci.1746-05.2005.

Luk, B., Mohammed, M., Liu, F. and Lee, F. J. S. (2015). A Physical Interaction between the Dopamine Transporter and DJ-1 Facilitates Increased Dopamine Reuptake. PLoS One. 10(8), e0136641. doi:10.1371/journal.pone.0136641.

Malicki, J. J. and Johnson, C. A. (2017). The Cilium: Cellular Antenna and Central Processing Unit. Trends Cell Biol. 27(2), 126–140. doi:10.1016/j.tcb.2016.08.002.

Massinen, S., Hokkanen, M. E., Matsson, H., Tammimies, K., Tapia-Páez, I., Dahlström-Heuser, V., Kuja-Panula, J., Burghoorn, J., Jeppsson, K. E., Swoboda, P., et al. (2011). Increased expression of the dyslexia candidate gene DCDC2 affects length and signaling of primary cilia in neurons. PloS One. 6(6), e20580. doi:10.1371/journal.pone.0020580.

Mitchison, H. M. and Valente, E. M. (2017). Motile and non-motile cilia in human pathology: from function to phenotypes. J. Pathol. 241(4), 564–564. doi:10.1002/path.4881.

Mizushima, N. and Levine, B. (2010). Autophagy in mammalian development and differentiation. Nat. Cell Biol. 12(9), 823–830. doi:10.1038/ncb0910-823.

Morgado, A. L., Xavier, J. M., Dionísio, P. A., Ribeiro, M. F. C., Dias, R. B., Sebastião, A. M., Solá, S. and Rodrigues, C. M. (2015). MicroRNA-34a Modulates Neural Stem Cell Differentiation by Regulating Expression of Synaptic and Autophagic Proteins. Mol. Neurobiol. 51(3), 1168–1183. doi:10.1007/s12035-014-8794-6.

Mykytyn, K., Nishimura, D. Y., Searby, C. C., Shastri, M., Yen, H.-J., Beck, J. S., Braun, T., Streb, L. M., Cornier, A. S., Cox, G. F., et al. (2002). Identification of the gene (BBS1) most commonly involved in Bardet-Biedl syndrome, a complex human obesity syndrome. Nat. Genet. 31(4), 435–438. doi:10.1038/ng935.

Nachury, M. V. and Mick, D. U. (2019). Establishing and regulating the composition of cilia for signal transduction. Nat. Rev. Mol. Cell Biol. 20(7), 389–405. doi:10.1038/s41580-019-0116-4.

Pertea, M., Pertea, G. M., Antonescu, C. M., Chang, T.-C., Mendell, J. T. and Salzberg, S. L. (2015). StringTie enables improved reconstruction of a transcriptome from RNA-seq reads. Nat. Biotechnol. 33(3), 290–295. doi:10.1038/nbt.3122.

Pevny, L. H. and Nicolis, S. K. (2010). Sox2 roles in neural stem cells. Int. J. Biochem. Cell Biol. 42(3), 421–424. doi:10.1016/j.biocel.2009.08.018.

Piasecki, B. P., Burghoorn, J. and Swoboda, P. (2010). Regulatory Factor X (RFX)-mediated transcriptional rewiring of ciliary genes in animals. Proc. Natl. Acad. Sci. U.S.A. 107(29), 12969–12974. doi:10.1073/pnas.0914241107.

Pierce, S. E., Tyson, T., Booms, A., Prahl, J. and Coetzee, G. A. (2018). Parkinsons disease genetic risk in a midbrain neuronal cell line. Neurobiol. Dis. 114, 53–64. doi:10.1016/j.nbd.2018.02.007.

Pitaval, A., Tseng, Q., Bornens, M. and Théry, M. (2010). Cell shape and contractility regulate ciliogenesis in cell cycle–arrested cells. J. Cell Biol. 191(2), 303–312. doi:10.1083/jcb.201004003.

Pletz, N., Medack, A., Rieß, E. M., Yang, K., Kazerouni, Z. B., Hüber, D. and Hoyer-Fender, S. (2013). Transcriptional activation of Odf2/Cenexin by cell cycle arrest and the stress activated signaling pathway (JNK pathway). Biochim. Biophys. Acta. 1833(6), 1338–1346. doi:10.1016/j.bbamcr.2013.02.023.

Reiter, J. F. and Leroux, M. R. (2017). Genes and molecular pathways underpinning ciliopathies. Nat. Rev. Mol. Cell Biol. 18(9), 533–547. doi:10.1038/nrm.2017.60.

Rohart, F., Gautier, B., Singh, A. and Cao, K.-A. L. (2017). mixOmics: An R package for ‘omics feature selection and multiple data integration. PLOS Comput. Biol. 13(11), e1005752. doi:10.1371/journal.pcbi.1005752.

Sarkisian, M. R. and Guadiana, S. M. (2015). Influences of Primary Cilia on Cortical Morphogenesis and Neuronal Subtype Maturation. Neuroscientist. 21(2), 136–151. doi:10.1177/1073858414531074.

Satija, R., Farrell, J. A., Gennert, D., Schier, A. F. and Regev, A. (2015). Spatial reconstruction of single-cell gene expression data. Nat. Biotechnol. 33(5), 495–502. doi:10.1038/nbt.3192.

Schildknecht, S., Pöltl, D., Nagel, D. M., Matt, F., Scholz, D., Lotharius, J., Schmieg, N., Salvo-Vargas, A. and Leist, M. (2009). Requirement of a dopaminergic neuronal phenotype for toxicity of low concentrations of 1-methyl-4-phenylpyridinium to human cells. Toxicol. Appl. Pharmacol. 241(1), 23–35. doi:10.1016/j.taap.2009.07.027.

Schneider, C. A., Rasband, W. S. and Eliceiri, K. W. (2012). NIH Image to ImageJ: 25 years of image analysis. Nat. Methods. 9(7), 671–675. doi:10.1038/nmeth.2089.

Scholz, D., Pöltl, D., Genewsky, A., Weng, M., Waldmann, T., Schildknecht, S. and Leist, M. (2011). Rapid, complete and large-scale generation of post-mitotic neurons from the human LUHMES cell line. J. Neurochem. 119(5), 957–971. doi:10.1111/j.1471-4159.2011.07255.x.

Senti, G. and Swoboda, P. (2008). Distinct isoforms of the RFX transcription factor DAF-19 regulate ciliogenesis and maintenance of synaptic activity. Mol. Biol. Cell. 19(12), 5517–5528. doi:10.1091/mbc.e08-04-0416.

Shah, R. R., Cholewa-Waclaw, J., Davies, F. C., Paton, K. M., Chaligne, R., Heard, E., Abbott, C. M. and Bird, A. P. (2016). Efficient and versatile CRISPR engineering of human neurons in culture to model neurological disorders. Wellcome Open Res. 1, 13. doi:10.12688/wellcomeopenres.10011.1.

Smirnova, L., Harris, G., Delp, J., Valadares, M., Pamies, D., Hogberg, H. T., Waldmann, T., Leist, M. and Hartung, T. (2016). A LUHMES 3D dopaminergic neuronal model for neurotoxicity testing allowing long-term exposure and cellular resilience analysis. Arch. Toxicol. 90(11), 2725–2743. doi:10.1007/s00204-015-1637-z.

Stanton, B. Z. and Peng, L. F. (2010). Small-molecule modulators of the Sonic Hedgehog signaling pathway. Mol. Biosys. 6(1), 44–54. doi:10.1039/b910196a.

Stepkowski, T. M, Wasyk, I., Grzelak, A. and Kruszewski, M. (2015). 6-OHDA-Induced Changes in Parkinson’s Disease-Related Gene Expression are not Affected by the Overexpression of PGAM5 in In Vitro Differentiated Embryonic Mesencephalic Cells. Cell. Mol. Neurobiol. 35(8), 1137–1147. doi:10.1007/s10571-015-0207-5.

Subramanian, A., Tamayo, P., Mootha, V. K., Mukherjee, S., Ebert, B. L., Gillette, M. A., Paulovich, A., Pomeroy, S. L., Golub, T. R. Lander, et al. (2005). Gene set enrichment analysis: A knowledge-based approach for interpreting genome-wide expression profiles. Proc. Natl. Acad. Sci. U.S.A. 102(43), 15545–15550. doi:10.1073/pnas.0506580102.

Sugiaman-Trapman, D., Vitezic, M., Jouhilahti, E.-M., Mathelier, A., Lauter, G., Misra, S., Daub, C. O., Kere, J. and Swoboda, P. (2018). Characterization of the human RFX transcription factor family by regulatory and target gene analysis. BMC Genomics. 19(1), 181. doi:10.1186/s12864-018-4564-6.

Tammimies, K., Bieder, A., Lauter, G., Sugiaman-Trapman, D., Torchet, R., Hokkanen, M. E., Burghoorn, J., Castrén, E., Kere, J., Tapia-Páez, I., et al. (2016). Ciliary dyslexia candidate genes DYX1C1 and DCDC2 are regulated by Regulatory Factor X (RFX) transcription factors through X-box promoter motifs. FASEB J. 30(10), 3578–3587. doi:10.1096/fj.201500124RR.

Thomas, S., Boutaud, L., Reilly, M. L. and Benmerah, A. (2019). Cilia in hereditary cerebral anomalies. Biol. Cell. 111(9), 217–231. doi:10.1111/boc.201900012.

Töhönen, V., Katayama, S., Vesterlund, L., Jouhilahti, E.-M., Sheikhi, M., Madissoon, E., Filippini-Cattaneo, G., Jaconi, M., Johnsson, A., Bürglin, T. R., et al. (2015). Novel PRD-like homeodomain transcription factors and retrotransposon elements in early human development. Nat. Commun. 6(1), 8207. doi:10.1038/ncomms9207.

van Dam, T. J., Wheway, G., Slaats, G. G., Huynen, M. A. and Giles, R. H. (2013). The SYSCILIA gold standard (SCGSv1) of known ciliary components and its applications within a systems biology consortium. Cilia. 2(1), 7. doi:10.1186/2046-2530-2-7.

Wagih, O. (2017). ggseqlogo: a versatile R package for drawing sequence logos. Bioinformatics. 33(22), 3645–3647. doi:10.1093/bioinformatics/btx469.

Wheway, G., Nazlamova, L. and Hancock, J. T. (2018). Signaling through the Primary Cilium. Front. Cell Dev. Biol. 6, 8. doi:10.3389/fcell.2018.00008

Wulle, I. and Schnitzer, J. (1989). Distribution and morphology of tyrosine hydroxylase-immunoreactive neurons in the developing mouse retina. Brain Res. Dev. Brain Res. 48(1), 59–72. doi:10.1016/0165-3806(89)90093-x.

Yan, J., Enge, M., Whitington, T., Dave, K., Liu, J., Sur, I., Schmierer, B., Jolma, A., Kivioja, T., Taipale, M., et al. (2013). Transcription Factor Binding in Human Cells Occurs in Dense Clusters Formed around Cohesin Anchor Sites. Cell. 154(4), 801–813. doi:10.1016/j.cell.2013.07.034.

Youn, Y. H. and Han, Y.-G. (2018). Primary Cilia in Brain Development and Diseases. Am. J. Pathol. 188(1), 11–22. doi:10.1016/j.ajpath.2017.08.031.

Zhang, S. and Cui, W. (2014). Sox2, a key factor in the regulation of pluripotency and neural differentiation. World J. Stem Cells. 6(3), 305–311. doi:10.4252/wjsc.v6.i3.305.

